# Duplexed CeTEAM drug biosensors reveal determinants of PARP inhibitor selectivity

**DOI:** 10.1101/2024.08.09.607390

**Authors:** Maria J. Pires, Alen Lovric, Seher Alam, Emanuele Fabbrizi, Dante Rotili, Mikael Altun, Nicholas C.K. Valerie

## Abstract

PARP inhibitors (PARPi) predominantly targeting PARP1 and PARP2 have revolutionized cancer therapy by selectively killing cancer cells with defective DNA repair. However, achieving PARP1 or PARP2-selective inhibitors is difficult due to their close structural homology. Selectivity profiling is typically done with purified proteins, but these lack the complexity of intracellular environments and could therefore be inaccurate. Here, we duplex PARP1 L713F-GFP and PARP2 L269A-mCherry CeTEAM drug biosensors to systematically characterize binding and cell cycle alterations of 27 PARPi at the single cell level. Our results reveal that most PARPi are generally equipotent for both PARPs, including the next-generation drug, senaparib. However, benzimidazole carboxamide (niraparib) derivatives demonstrated PARP1-selective tendencies, while pthalazinones (olaparib) favored PARP2. AZD5305, a reported PARP1-selective inhibitor with characteristics of both series, was the exception and appears ∼1600-fold more potent towards PARP1. In agreement with current understanding, we see that PARP trapping phenotypes positively correlate with PARP1/2 binding potency, while some potent binders, such as veliparib, did not – likely reflecting their allosteric influence on DNA retention. We also assessed the effect of the PARP1/2 active site component, HPF1, on intracellular PARPi binding and see that HPF1 depletion elicits slight deviations in apparent binding potency, while contributing additively to PARP-DNA trapping phenotypes. The PARP1/2 CeTEAM platform thus provides a structural roadmap for the development of selective PARPi and should facilitate the discovery of better targeted therapies. Furthermore, our results highlight that multiplexing CeTEAM biosensors and layered genetic perturbations can systematically profile determinants of intracellular drug selectivity.

## Introduction

PARP inhibitors (PARPi) have revolutionized treatment for cancers characterized by deficiencies in DNA repair mechanisms, particularly those involving mutations to the homologous recombination repair proteins, BRCA1 and BRCA2 (1). These drugs preferentially block the ability of PARP1 and PARP2 to initiate repair of DNA damage and selectively kill cancer cells that are unable to repair double strand breaks (DSBs) based on the concept of synthetic lethality (2, 3). This effect is further driven by the ability of some PARPi to trap PARP1 and PARP2 onto DNA, thereby creating more irreparable DNA damage (1, 4).

Most PARPi, consisting of a nicotinamide mimetic pharmacophore, indiscriminately target both PARP1 and PARP2 and, to a lesser extent, other PARP paralogs (4). PARP1 and PARP2 have substantially different N-terminal DNA binding domains but share homologous catalytic domains and allosteric regulatory mechanisms (5, 6). Therefore, specific targeting of their enzymatic activities has been challenging to date, as the structural variation in current PARPi conveys similar inhibition of both paralogs but very different allosteric outcomes (5, 7). Nonetheless, each has unique biological functions warranting preferential targeting (8), which could convey clinical ramifications but also create therapeutic opportunities. For example, the hematological toxicity of PARPi has been attributed to an on-target, adverse effect of PARP2 inhibition due to its essential role in erythroid progenitor differentiation (9, 10). Thus, despite an abundance of potent, clinical-grade inhibitors, the need for truly selective, next generation PARPi remains high.

Small molecule selectivity is traditionally assessed with purified proteins *in vitro*, as more exact calculations of the inhibitory potency and characterizations of the binding modality can be performed. However, selectivity profiling in cellular contexts, as opposed to with purified proteins, has unveiled disparities in inhibitor performance, suggesting that the physiological, intracellular environment can significantly influence the behavior of inhibitors (11). For example, the interaction of PARP1 and PARP2 with co-factors like Histone PARylation Factor 1 (HPF1) can further complicate the pharmacodynamics of PARPi (12). HPF1 modulates the PARylation activity of PARP1 and PARP2 by effectively completing their enzymatic pockets, and, in turn, tuning their ability to interact with and modify histones during chromatin remodeling and DNA repair (12, 13). Thus, it is reasonable to expect HPF1 to influence PARPi binding and activity. Indeed, early indications suggest that HPF1 can modulate PARPi binding potency *in vitro* (14, 15) and PARPi sensitivity *in cellulo* (16), but the effect on PARPi binding in cells has not been elucidated.

Here, we utilize the recently developed cellular target engagement by accumulation of mutants (CeTEAM) assay (17) to comprehensively profile intracellular selectivity of PARPi towards PARP1 and PARP2. CeTEAM leverages stability-dependent biosensors that can be multiplexed with other readouts of downstream pharmacology to follow drug-target interactions *in cellulo*. By means of a duplexed, dual fluorescence CeTEAM platform, we compare the binding dynamics of reported PARPi for PARP1 and PARP2 drug biosensors and integrate ensuing cell cycle changes to generate a compendium of PARP1/2 binding and trapping profiles in live cells. While most PARPi equipotently engage PARP1/2, many benzimidazole carboxamide (niraparib) derivatives possess slight selectivity for PARP1 (>10-fold), while pthalazinones (olaparib) favor PARP2 binding. However, AZD5305 displays ∼1600-fold selectivity towards PARP1, thus representing a truly selective PARPi. Furthermore, PARP1/2 binding potency has strong positive correlation with PARP trapping phenotypes with the notable exception of veliparib and venadaparib, which may reflect their allosteric tendencies. We also assess the influence of HPF1 on these readouts and find that HPF1 knockdown modestly affects PARP1/2 binding but additively promotes PARP trapping phenotypes. Our findings reveal potential structural determinants of PARPi selectivity towards PARP1 or PARP2 and provide an avenue for integrating genetic screens into cell-based drug binding and selectivity assays. The multiplexed CeTEAM selectivity platform should be a valuable tool for evaluating the PARP1/2 binding and trapping of next-generation PARP inhibitors.

## Results

### Validation of a duplexed PARP1/2 CeTEAM fluorescent biosensor system

Despite almost all clinical PARPi targeting both PARP1 and PARP2, there is a paucity of data regarding intracellular PARP2 drug-target engagement, especially regarding relative potency (18–20). In our hands, we were unable to detect discernible shifts in PARP2 stability by the cellular thermal shift assay (CETSA), despite seeing expected increases in soluble PARP1 upon olaparib addition (21, 22), which may partially explain the limited data on PARP2 engagement in cells (**Supplementary Fig. 1A-C**). We previously demonstrated the CeTEAM amenability of GFP-tagged PARP1 L713F and the analogous mutant in PARP2 (L269A) individually with multiple PARPi by comprehensive dose-response profiling (17). To build a truly multiplexed platform for PARPi selectivity profiling, we utilized spectrally distinct fluorescent tags for simultaneous monitoring of PARP1 L713F (GFP) and PARP2 L269A (mCherry) within the same cellular context (**Fig. 1**). We validated this system by western blot analysis after a 24-hour PARPi treatment using talazoparib, olaparib, niraparib, veliparib, and 3-aminobenzamide (3-AB), with iniparib as a negative control (**Fig. 1A and B**). All PARPi showed dose-dependent stabilization of PARP1 and PARP2 except for iniparib. As before (17), 3-AB required higher concentrations for similar engagement. Unexpectedly, niraparib stabilized PARP2 L269A significantly worse than PARP1 L713F at 1 µM.

**Figure 1.**
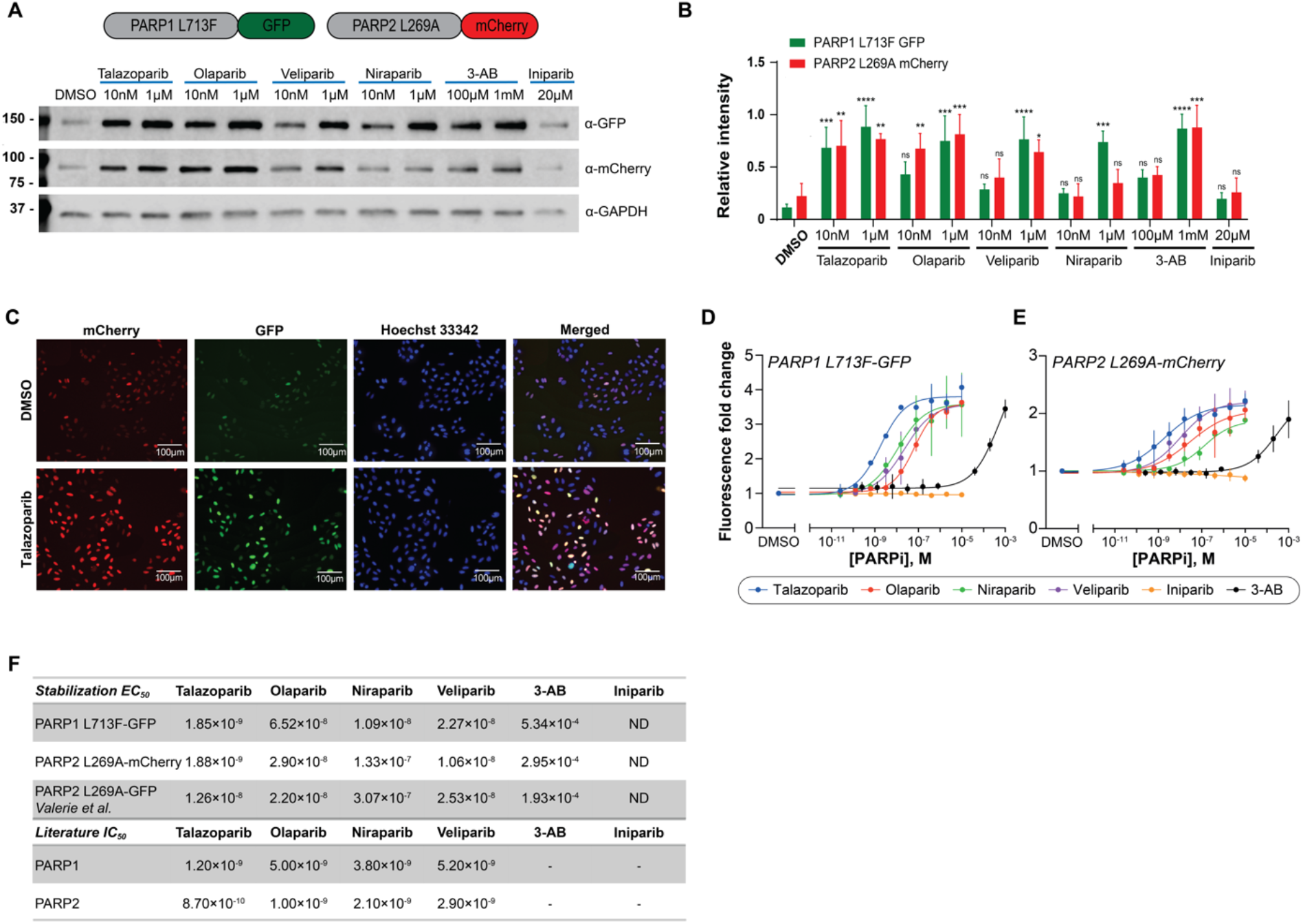
Establishment of a duplexed PARP1/2 CeTEAM drug biosensor platform. **A.** A representative western blot of PARP1 L713F-GFP and PARP2 L269A-mCherry abundance after 24-hour treatment with PARP inhibitors (PARPi) at indicated concentrations. **B.** Densitometric quantification of PARP1 L713F-GFP and PARP2 L269A-mCherry relative to GAPDH and normalized to the highest value observed for each reporter. Means shown ± SD (n=3). Statistical significance indicated as not significant (ns), * – *p* < 0.05, ** – *p* < 0.01, *** – *p* < 0.001, and **** – *p* < 0.0001 as determined by one-way ANOVA with Dunnett’s post-test comparing each condition to the DMSO vehicle control. **C.** Representative live-cell fluorescence micrographs of L713F-GFP (green) and L269A-mCherry (red) treated with DMSO or 10 µM talazoparib for 24 hours. Cell nuclei identified by Hoechst staining (blue) and scale bar = 100µm. **D.** Live-cell L713F-GFP fold change by fluorescence microscopy following a concentration gradient with the indicated PARPi for 24 hours and normalization to DMSO controls. Means ± SD (n=3) with lines-of-best-fit shown. **E.** Live-cell L269A-mCherry fold change by fluorescence microscopy, as in **D**. Means ± SD (n=3) with lines-of-best-fit shown. **F.** Observed stabilization EC_50_ values for PARP1 L713F-GFP and PARP2 L269A-mCherry with indicated PARPi compared with PARP2 L269A-GFP data from *Valerie et al.* Reported biochemical IC_50_ values given for reference. ND – not determined.

Live-cell fluorescence microscopy confirmed these results, demonstrating essentially equipotent stabilization EC_50_ values with both biosensors, which ranged from ∼2 nM (talazoparib) to ∼300-500 µM (3-AB). These values were generally in close agreement with reported biochemical IC_50_ and K_i_ values (**Fig. 1C-F**). Notably, however, niraparib binding to PARP2 L269A-mCherry was worse by >10-fold (∼11 nM *vs* ∼133 nM) (23). Findings with L269A-mCherry were consistent with earlier stabilization EC_50_ values from PARP2 L269A-GFP (17), suggesting that the choice of fluorescent protein tag did not specifically contribute to observed stabilization changes (**Fig. 1F**). Thus, our PARP1/2 duplexed CeTEAM biosensor system is a capable platform for detailed pharmacological assessments of PARP inhibitors.

### Re-screening of PARP1 L713F-nLuc stabilizers with the PARP1/2 CeTEAM assay

We previously screened a ∼1200 compound drug-like library for PARP1 L713F biophysical perturbagens and found >90% of potent PARP1i, as well as several non-PARPi that also stabilized the nLuc biosensor (17). To determine if these molecules also affected PARP2 L269A stability, we then re-screened the original hits with the duplexed PARP1/2 CeTEAM assay (**Fig. 2A**). This also afforded the opportunity to validate PARP1 L713F stabilization by an orthogonal readout and integrate cell cycle measurements in response to drug treatment, which could further inform on differential pharmacology. In total, the re-screen contained the annotated PARPi and 14 non-PARPi tested previously, as well as two additional PARPi reported to be PARP1-(MC2050) and PARP2-selective (MC3474; Compound 1), respectively (24). The compounds were tested at 10 µM and biosensor abundance was measured after 16 hours. Hits were defined as having statistically significant (*p* < 0.05) abundance changes from the DMSO (negative) control.

**Figure 2.**
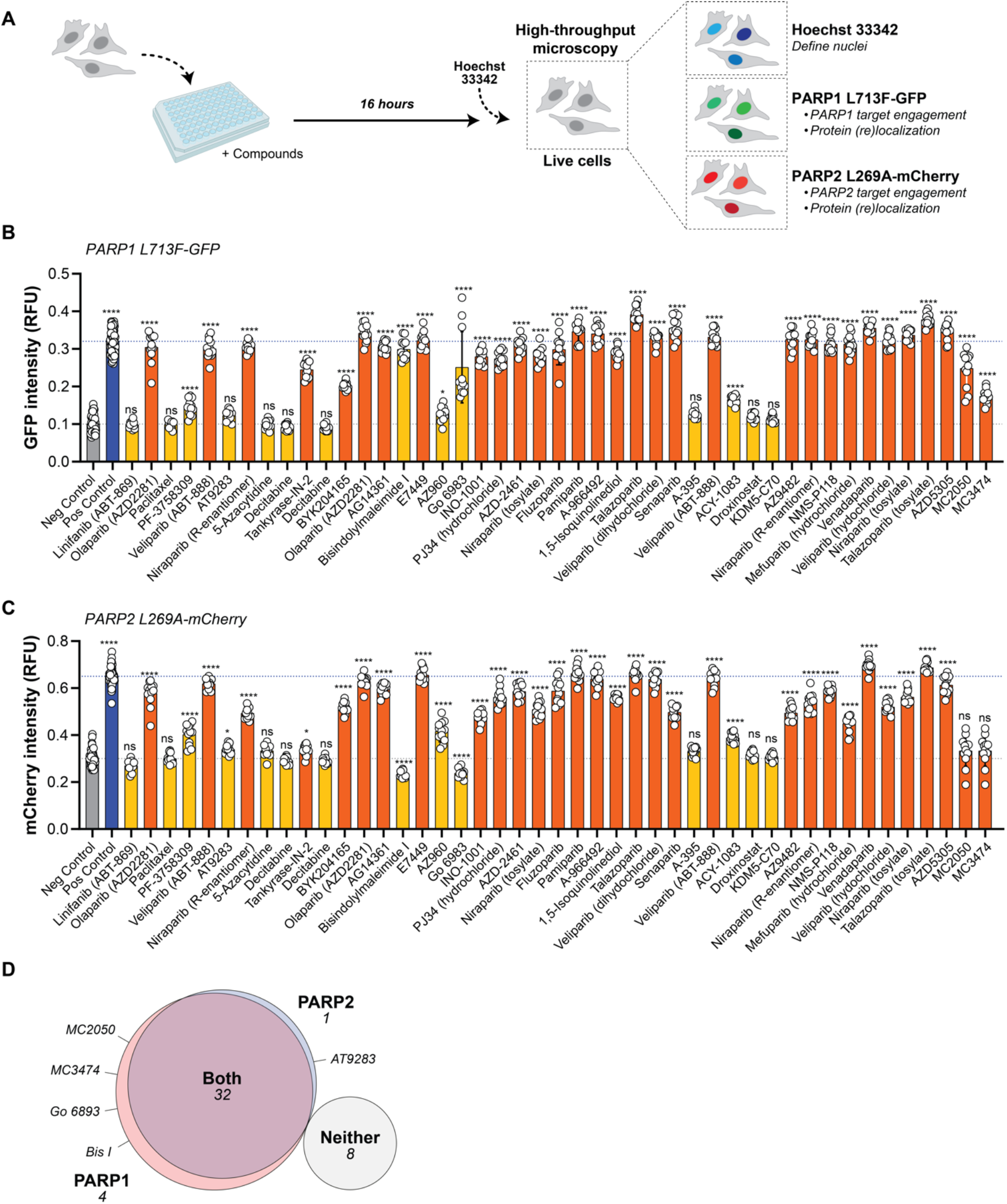
Rescreening of potential PARP1/2 biophysical modulators with duplexed PARPi biosensors. **A.** Overview of rescreening procedure for PARP1 (GFP)/PARP2 (mCherry) target engagement by live-cell fluorescent microscopy. Nuclei are identified by Hoechst 33342 staining. **B.** PARP1 L713F-GFP intensity following 10 µM PARPi (orange, 31 total) and non-PARPi (yellow, 14 total). Negative (DMSO, grey) and positive controls (10 µM veliparib, blue) are shown with intensity reference lines. Means shown for three wells (four images per well) ± SD. Statistical significance denoted as not significant (ns), * – *p* < 0.05, ** – *p* < 0.01, *** – *p* < 0.001, and **** – *p* < 0.0001 as determined by one-way ANOVA with Dunnett’s post-test comparing each condition to the DMSO vehicle control. **C.** PARP2 L269A-mCherry intensities for the rescreened compounds, as in **B**. **D.** Venn diagram representation of statistically significant hits for PARP1 and PARP2. Compounds affecting neither target are also shown, while those stabilizing only one target are named. RFU – relative fluorescence units.

All PARPi significantly increased PARP1 L713F-GFP abundance, but only a few non-PARPi (PF-3758309, bisindolylmaleimide I, AZ960, Go 6983, and ACY-1083) also met this threshold, essentially mirroring the L713F-nLuc biosensor reported earlier (17). Notably, the DNMT trappers, decitabine and azacitidine, did not meaningfully increase L713F-GFP, in contrast to the nLuc reporter reported earlier (17). Almost all PARPi also significantly increased PARP2 L269A-mCherry abundance, except for MC2050 and MC3474 (**Fig. 2C**). AZD5305, a compound previously identified as a highly specific PARP1 inhibitor (25, 26), comparably stabilized PARP2 at this high dose (**Fig. 2B and C**). The same non-PARPi that increased L713F-GFP also affected PARP2 L269A-mCherry, except for the two protein kinase C inhibitors (PKCi; bisindolylmaleimide I and Go 6983), which, in contrast to PARP1 L713F, significantly decreased PARP2 L269A abundance. Another non-PARPi, AT9283, showed a modest but significant effect on PARP2 that was not seen with PARP1.

We then performed a cross-comparison of hits for PARP1 and PARP2 to help identify any clear trends (**Fig. 2D**). Of the compounds tested, 32 significantly stabilized both PARP1 and PARP2, while 8 did not appreciably affect either. All non-responsive molecules were annotated non-PARPi. Therefore, apart from select molecules, effects on PARP2 L269A abundance generally aligned with PARP1 L713F at the 10 µM screening concentration.

### Intracellular selectivity profiling of PARPi with CeTEAM

Initial follow-up of non-PARPi hits confirmed that the broad-spectrum PKC inhibitors, bisindolylmaleimide I and Go 6983 selectively increased PARP1 L713F-GFP abundance in a dose-responsive manner (**Supplementary Fig. 2A-B**). However, systematic dissection of these molecules revealed that both are autofluorescence artifacts at emission wavelengths ∼600 nm (**Supplementary Fig. 3A-E**), in agreement with earlier reports using fluorescence-based detection systems (27).

In parallel with our analysis of non-PARPi, we conducted dose-response experiments with the expanded PARPi library using the duplexed PARP1/2 biosensors to establish a compendium of PARP1 and PARP2 binding in live cells (**Fig. 3A**). Comparative analysis of the dose-responses for PARP1 L713F-GFP (**Fig. 3B**) and PARP2 L269A-mCherry (**Fig. 3C**), alongside literature-derived IC_50_ values, facilitated a robust evaluation of each inhibitor’s cellular efficacy, as summarized in **Table 1** and depicted in **Supplementary Figure 4**. As before, talazoparib, olaparib, veliparib, and 3-AB were equipotent for PARP1 and PARP2, while iniparib was inactive and niraparib was ∼14-fold more potent towards PARP1. Unsurprisingly, clinical-grade PARPi were typically the most potent for both PARPs (low nanomolar), while early generation PARPi (*e.g.*, PJ34, BYK204165, 1,5-isoquinolinediol, 3-AB) or those not intended to directly target PARP1/2 (*e.g.*, Tankyrase-IN-2, ME0328) were significantly worse (**Table 1**). Tankyrase-IN-2 is reported to predominantly target TNKS1/2 but has low level activity towards PARP1(28), while ME0328 preferentially targets PARP3 (29). Furthermore, the next generation PARPis, senaparib (IMP4297) and venadaparib, Click or tap here to enter text.equipotently engaged PARP1/2 biosensors in the low nanomolar range like other potent PARPi, such as A-966492 and rucaparib (**Table 1**), in line with biochemical inhibition data towards PARP1 and PARP2 (senaparib: 0.48 and 1.6 nM, venadaparib: 0.8 and 3.0 nM, respectively) (31, 32).Click or tap here to enter text.

**Figure 3.**
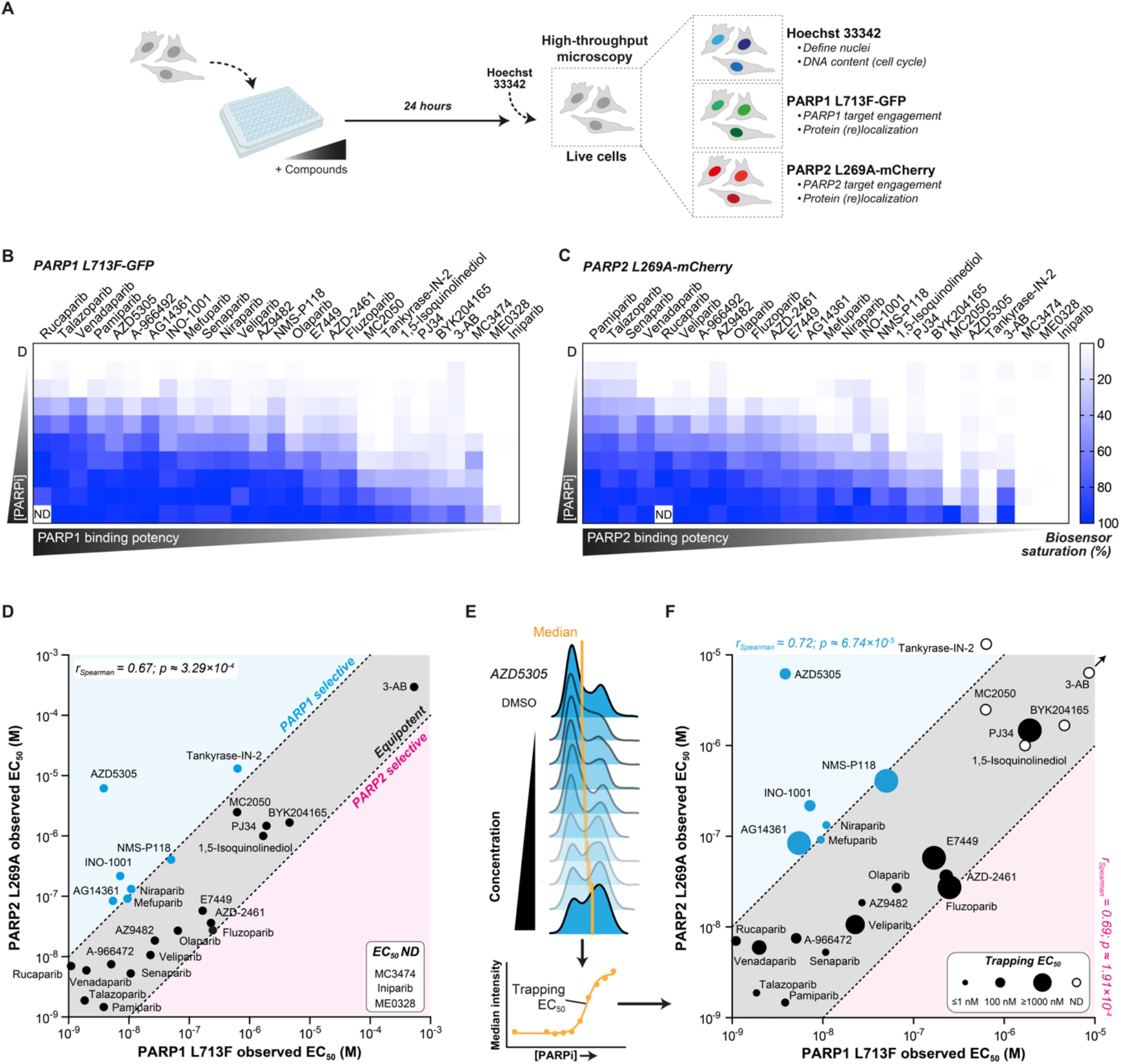
Intracellular PARP inhibitor selectivity and trapping profiling with CeTEAM. **A.** Experimental procedure for dose-dependent PARP1 L713F-GFP and PARP2 L269A-mCherry stabilization and cell cycle perturbations (Hoechst 33342) by PARPi with live-cell fluorescent microscopy. **B.** Relative PARP1 L713F-GFP intensity expressed as fold change DMSO control after 24 hours of a PARPi gradient. Means shown from n=3. **C.** Relative PARP2 L269A-mCherry intensity expressed as fold change DMSO control after 24 hours of a PARPi gradient. Means shown from n=3. **D.** Two-dimensional comparison of PARPi stabilization EC_50_ values towards PARP1 L713F-GFP and PARP2 L269A-mCherry. Blue – ≥10-fold PARP selectivity (PARP1 selective); red – ≥10-fold PARP2 selectivity (PARP2 selective); grey – <10-fold selectivity difference (equipotent). Spearman correlation r = 0.67; p ≈ 3.29×10^-4^. Values derived from n=3 replicates from **B** and **C**. **E.** Derivation of PARP trapping EC_50_ from the median Hoechst intensity (DNA content). **F.** Three-dimensional comparison of PARP1 (x-axis; blue) and PARP2 (y-axis; red) engagement EC_50_ values and PARP trapping EC_50_ values (circle size) for tested PARPi. Circle size – ≤1 nM (smallest) to ≥1000 nM (largest), or open circle – EC_50_ not determined (ND). A representative experiment is used for trapping EC_50_ overlay. Spearman correlation r = 0.72; p ≈ 6.74×10^-5^ for PARP1. Spearman correlation r = 0.69; p ≈ 1.91×10^-4^ for PARP2.

**Table 1.** Comparison of CeTEAM stabilization EC_50_ values and reported biochemical IC_50_ values of tested PARPi towards PARP1/2 (M)

| *PARPi* | PARP1 IC<sub>50</sub> | PARP1 L713F-GFP EC<sub>50</sub> | PARP2 IC<sub>50</sub> | PARP2 L269A-mCherry EC<sub>50</sub> |
| --- | --- | --- | --- | --- |
| *Talazoparib* | <sup>a</sup>1.20×10<sup>-9</sup> | 1.854×10<sup>-9</sup> | <sup>a</sup>8.70×10<sup>-10</sup> | 1.875×10<sup>-9</sup> |
| *Pamiparib* | <sup>a</sup>9.00×10<sup>-10</sup> | 3.839×10<sup>-9</sup> | <sup>a</sup>5.00×10<sup>-10</sup> | 1.467×10<sup>-9</sup> |
| *Niraparib* | <sup>a</sup>3.80×10<sup>-9</sup> | 1.093×10<sup>-8</sup> | <sup>a</sup>2.10×10<sup>-9</sup> | 1.325×10<sup>-7</sup> |
| *Veliparib* | <sup>a</sup>5.20×10<sup>-9</sup> | 2.274×10<sup>-8</sup> | <sup>a</sup>2.90×10<sup>-9</sup> | 1.062×10<sup>-8</sup> |
| *Rucaparib* | <sup>a</sup>1.40×10<sup>-9</sup> | 1.110×10<sup>-9</sup> | <sup>a</sup>1.40×10<sup>-9</sup> | 7.013×10<sup>-9</sup> |
| *A-966492* | <sup>b</sup>1.00×10<sup>-9</sup> | 5.081×10<sup>-9</sup> | <sup>b</sup>1.50×10<sup>-9</sup> | 1.201×10<sup>-8</sup> |
| *Olaparib* | <sup>a</sup>5.00×10<sup>-9</sup> | 6.520×10<sup>-8</sup> | <sup>a</sup>1.00×10<sup>-9</sup> | 2.689×10<sup>-8</sup> |
| *PJ34* | <sup>a</sup>1.10×10<sup>-7</sup> | 1.913×10<sup>-6</sup> | <sup>a</sup>8.60×10<sup>-8</sup> | 1.471×10<sup>-6</sup> |
| *Mefuparib* | <sup>a</sup>3.20×10<sup>-9</sup> | 9.477×10<sup>-9</sup> | <sup>a</sup>1.90×10<sup>-9</sup> | 9.156×10<sup>-8</sup> |
| *MC3474* | <sup>a</sup>2.04×10<sup>-6</sup> | ND | <sup>a</sup>3.4×10<sup>-7</sup> | ND |
| *MC2050* | <sup>a</sup>9.30×10<sup>-9</sup> | 6.246×10<sup>-7</sup> | <sup>a</sup>5.49×10<sup>-7</sup> | 2.496×10<sup>-6</sup> |
| *AZD5305* | <sup>a</sup>3.00×10<sup>-9</sup> | 3.847×10<sup>-9</sup> | <sup>a</sup>1.40×10<sup>-6</sup> | 6.172×10<sup>-6</sup> |
| *Tankyrase-IN-2* | <sup>a</sup>7.1×10<sup>-7</sup> | 7.885×10<sup>-7</sup> | - | 1.315×10<sup>-5</sup> |
| *AZ9482* | <sup>b</sup>1.00×10<sup>-9</sup> | 2.701×10<sup>-8</sup> | <sup>b</sup>1.00×10<sup>-9</sup> | 1.849×10<sup>-8</sup> |
| *Senaparib* | <sup>a</sup>4.80×10<sup>-10</sup> | 1.074×10<sup>-8</sup> | <sup>a</sup>1.60×10<sup>-9</sup> | 5.257×10<sup>-9</sup> |
| *INO-1001* | <sup>b</sup><5.0×10<sup>-8</sup> | 7.198×10<sup>-9</sup> | - | 2.172×10<sup>-7</sup> |
| *NMS-P118* | <sup>a</sup>9.00×10<sup>-9</sup> | 5.016×10<sup>-8</sup> | <sup>a</sup>1.39×10<sup>-6</sup> | 4.076×10<sup>-7</sup> |
| *Venadaparib* | <sup>a</sup>1.40×10<sup>-9</sup> | 1.986×10<sup>-9</sup> | <sup>a</sup>1.00×10<sup>-9</sup> | 5.935×10<sup>-9</sup> |
| *AG14361* | <sup>b</sup>1.40×10<sup>-8</sup> | 5.458×10<sup>-9</sup> | - | 8.399×10<sup>-8</sup> |
| *E7449* | <sup>a</sup>2.00×10<sup>-9</sup> | 1.679×10<sup>-7</sup> | <sup>a</sup>1.00×10<sup>-9</sup> | 5.776×10<sup>-8</sup> |
| *AZD2461* | <sup>a</sup>5.00×10<sup>-9</sup> | 2.294×10<sup>-7</sup> | <sup>a</sup>2.00×10<sup>-9</sup> | 3.626×10<sup>-8</sup> |
| *Fluzoparib* | <sup>a</sup>1.46×10<sup>-9</sup> | 2.49×10<sup>-7</sup> | - | 2.734×10<sup>-8</sup> |
| *BYK204165* | <sup>a</sup>7.35×10<sup>-9</sup> | 4.607×10<sup>-6</sup> | <sup>a</sup>4.17×10<sup>-6</sup> | 1.669×10<sup>-6</sup> |
| *1,5-Isoquinolinediol* | <sup>b</sup>1.8-3.7×10<sup>-7</sup> | 1.694×10<sup>-6</sup> | - | 1.001×10<sup>-6</sup> |
| *ME0328* | <sup>a</sup>6.30×10<sup>-6</sup> | ND | <sup>a</sup>1.08×10<sup>-5</sup> | ND |
| *3-AB* | <sup>b</sup>5.00×10<sup>-8</sup> | 5.338×10<sup>-4</sup> | - | 2.953×10<sup>-4</sup> |
| *Iniparib* | - | ND | - | ND |

<sup>a</sup> Biochemical assay; <sup>b</sup> Cell-based assay; ND – not determined

Some PARPi also bound PARP1 and PARP2 much worse than anticipated. For example, PJ34 was >10-fold worse towards both PARPs in our assay as compared to reported IC_50_ values (**Fig. 3D and Table 1**). Additionally, despite published specificity for PARP2 (24), MC3474 was one of the least potent PARPi in our assay, which may relate to its relatively high biochemical IC_50_ values (**Supplementary Fig. 4 and Table 1**). These demonstrated a similar phenomenon to other early PARPi (*e.g.*, 3-AB), where intracellular potency is significantly compromised.

Most of the expanded PARPi library were essentially equipotent binders of both PARPs, but some appeared to demonstrate slight selectivity towards PARP1 or PARP2. To facilitate comparisons of PARPi selectivity, we performed a pairwise analysis of PARP1 and PARP2 biosensor saturation for each PARPi and compared their apparent EC_50_ values (**Fig. 3D**). We arbitrarily defined selectivity as > ∼10-fold differential potency (cyan or magenta regions) and equipotency as <10-fold preference (grey region) for either PARP. As expected, most inhibitors fell within the equipotency designation, as supported by PARP1 and PARP2 biosensor binding potencies being highly correlative (r_Spearman_ = 0.67, *p* ≈ 0.000329).

While some inhibitors had negligible selectivity towards PARP1 (*e.g.*, rucaparib and MC2050), AZD5305, INO-1001, niraparib, AG14361, mefuparib, NMS-P118, and Tankyrase-IN-2 definitively favored PARP1 binding, although Tankyrase-IN-2 was significantly less potent overall. At 10 µM, we initially saw that AZD5305 exhibited comparable stabilization of both biosensors (**Fig. 2B and C**).

However, it displayed markedly higher specificity towards PARP1 over PARP2 by extended dose-response, as evidenced by a marked difference in stabilization EC_50_ values – 3.9 nM *vs* 6.2 µM, respectively (∼1600-fold; **Table 1**, **Fig. 3D**). Our measures of apparent PARP1/2 binding for AZD5305 align well with reported biochemical and intracellular IC_50_ values (25, 26). We could also confirm the slight specificity of MC2050 (24) for PARP1 in our cellular assays (PARP1: 625 nM, PARP2: 2482 nM), albeit with significantly worse efficacy than AZD5305 despite their similar biochemical potencies. Prior to the disclosure of AZD5305, NMS-P118 was reported as one of the first selective PARP1i (∼150-fold over PARP2, (31). However, our results reflected a much smaller selectivity window in cells (∼10-fold; 50.2 and 408 nM, respectively). Interestingly, the early generation PARPi, INO-1001, was quite robust in our assay and yielded a surprising ∼30-fold preference for PARP1 over PARP2 (apparent PARP1 EC_50_: ∼7.9 nM, PARP2 EC_50_: ∼217.2 nM), although we did not find reports with conclusive data describing its PARP1/2 inhibitory potency.

While PARP1-selectice molecules have been disclosed, to the best of our knowledge, there are no truly PARP2-selective inhibitors. In our assay, AZD2461 and fluzoparib demonstrated slight, but insignificant, selectivity for PARP2 (∼6- and 9-fold, respectively; **Table 1**, **Fig. 3D**). Of the remaining molecules, none demonstrated appreciable preference for PARP2 binding.

In addition to live-cell fluorescence microscopy, we validated these findings for many PARPi by western blot analysis – confirming that detected fluorescence changes were proportional to protein abundance changes (**Supplementary Fig. 5A and B**). Collectively, these results suggest that a duplexed PARP1/2 CeTEAM platform can effectively profile PARPi selectivity in a scalable assay with live, single cells.

### PARP trapping phenotypes strongly correlate with PARP1/2 binding potency

In addition to the synthetic lethal DNA damage elicited in homologous recombination-defective cancer cells, many clinical PARPi can trap PARP1 and PARP2 on DNA as part of their anticancer mechanism of action (1, 32). It has previously been shown that measuring DNA content (cell cycle) is a suitable readout of the DNA replication stress/S-phase delay and subsequent G2 arrest resulting from PARP trapping (33). Similarly, we have also utilized the cell permeable stain, Hoechst 33342, as a proxy to profile trapping dynamics in live cells with the PARP1 L713F-GFP CeTEAM assay (17). We defined PARP trapping as PARPi that both engaged the PARP1 biosensor and elicited an S/G2 shift in the cell cycle. To profile how this expanded library of PARPi influenced cell cycle dynamics in relation to PARP1 and PARP2 binding, our PARPi library dose-response experiments in PARP1/2 biosensor cells were complemented with Hoechst 33342 (**Fig. 3A, Supplementary Fig. 6**). As before, this method enabled us to generate crude estimations of the cell cycle and estimated DNA trapping EC_50_ values by shifts in median Hoechst intensity (**Fig. 3E**, **Table 2**). In this case, a lower EC_50_ indicates more effective PARPi trapping, whereas a higher EC_50_ suggests reduced trapping. Further, juxtaposing this data with PARP1/2 biosensor binding can give insights to potential on- *vs* off-target activity of PARPi.

**Table 2.** DNA trapping EC_50_ values of tested PARPi from median Hoechst intensity changes.

| *PARPi* | DNA trapping EC<sub>50</sub> (M) |
| --- | --- |
| *Talazoparib* | 2.78e-010 |
| *Pamiparib* | 2.96e-009 |
| *Niraparib* | 2.85e-009 |
| *Veliparib* | 5.28e-007 |
| *Rucaparib* | 2.37e-008 |
| *A-966492* | 4.16e-008 |
| *Olaparib* | 2.57e-008 |
| *PJ34* | 3.75e-006 |
| *Mefuparib* | 2.36e-009 |
| *MC3474* | ND |
| *MC2050* | ND |
| *AZD5305* | 6.07e-008 |
| *Tankyrase-IN-2* | ND |
| *AZ9482* | 3.51e-010 |
| *Senaparib* | 3.98e-010 |
| *INO-1001* | 5.97e-008 |
| *NMS-P118* | 5.88e-006 |
| *Venadaparib* | 1.97e-007 |
| *AG14361* | 1.12e-006 |
| *E7449* | 3.97e-006 |
| *AZD2461* | 1.46e-007 |
| *Fluzoparib* | 4.63e-005 |
| *BYK204165* | ND |
| *1,5-Isoquinolinediol* | ND |
| *ME0328* | ND |
| *3-AB* | ND |
| *Iniparib* | ND |

ND – not determined

Our scaled approach also afforded the possibility of gauging the quality of PARP trapping phenotypes based on concentration-dependent cell cycle dynamics (**Supplementary Fig. 6**). Visual inspection of the cell cycle distributions revealed that trapping phenotypes do not manifest uniformly. For example, talazoparib, a potent PARP1/2 trapper, exhibited an initial increase in S/G2-phase cells at low doses but transitioned to primarily an S-phase delay at higher doses. This is consistent with a tolerable level of PARP retention-related DNA damage at low doses that permits continuation through S-phase and then triggers the G2-phase DNA damage checkpoint, while at higher doses, the extent of DNA replication-associated damage prohibits completion of S-phase. Senaparib also demonstrated such a profile, in support of it being a potent PARP1/2 binder and trapper. On the other hand, more modestly trapping PARPi, such as venadaparib or AZD5305, induce G2-phase accumulation but fail to progress to predominantly S-phase arrest at higher concentrations, perhaps due to less efficient PARP retention by these inhibitors.

We then combined the median Hoechst intensity changes (trapping EC_50_ values) with the PARP1 L713F-GFP and PARP2 L269A-mCherry to give insights on how trapping may be influenced by PARP1 and/or PARP2 binding. Overall, inhibitors that were the most potent binders of both PARP1 and PARP2 generally yielded the most potent trapping phenotypes, whereas the less potent molecules were generally poor at generating these phenotypes (**Fig. 3F**, **Table 2**). Both PARP1 and PARP2 biosensor binding potency highly correlated with PARP trapping phenotypes (r_Spearman_ = 0.72 [*p* ≈ 0.0000674] and 0.69 [*p* ≈ 0.000191], respectively). Among these, the next generation PARPi, senaparib, was among the most potent elicitors of PARP trapping phenotypes (0.40 nM trapping EC_50_), along with talazoparib and AZ9482 (0.28 and 0.35 nM, respectively). There were some exceptions to this trend – veliparib was a potent binder to both PARP1 and PARP2 but was exceptionally worse at trapping PARP to DNA (528 nM trapping EC_50_), in line with previous data (17, 32). Similarly, venadaparib potently engaged both PARP1 and -2 in our assay (1.99 and 5.94 nM, respectively) but had a corresponding trapping character EC_50_ value of 197 nM. The PARP1-selective inhibitor, AZD5305 also demonstrated a robust trapping phenotype but was comparatively inferior to other potent, dual PARP1/2 inhibitors, such as senaparib (60.7 nM *vs* 0.4 nM trapping EC_50_, respectively). On the other hand, PJ34, Tankyrase- IN-2, and 3-AB, which were among the worst PARP1/2 binders, had minimal (PJ34 – 3.75 µM EC_50_) or no observable trapping activity (Tankyrase-IN-2, 3-AB). Thus, our observations align with the prevailing view that trapping is predominantly driven by inhibitory potency but can also be strongly influenced by allosteric effects.

### Differential PARP1/2 selectivity between two primary PARPi pharmacophores

Apart from a handful of reported PARPi, most clinical drugs equipotently inhibit PARP1/2 activity (4), but enzymatic IC_50_ values may not necessarily align with biophysical measures of binding. Insights from biophysical readouts can inform on how well a compound binds to the target (34), which may be even more informative in a physiologically relevant, intracellular environment. From the literature, we note that niraparib is reported to inhibit PARP1 and PARP2 activity equipotently (3.8 and 2.1 nM, respectively (35)); however, biophysical measures have indicated that niraparib binds ∼50-fold worse to PARP2. We noticed similar biophysical trends for other benzimidazole carboxamide derivatives, specifically those with similar structures to niraparib (rucaparib, veliparib, NMS-P118), while other structural classes of PARPi (olaparib, talazoparib) more closely resembled the equipotency of enzymatic assays. We then revisited our PARP1/2 biosensor binding data and saw that the PARPi with slight PARP1 selectivity were enriched for these same inhibitor subtypes – namely AG14361, mefuparib, niraparib, and NMS-P118 (**Fig. 3D**). Likewise, the molecules with modest PARP2 selectivity appeared to be enriched with pthalazinone derivatives similar to olaparib. These insights suggested that PARP1 and PARP2 selectivity may already be engrained in prevailing PARPi pharmacophores.

To investigate these observations more systematically, we employed the cheminformatics server from ChemMine Tools to perform binned clustering and multidimensional scaling (2D MDS (36)) on the PARPi library. These algorithms consider atom-by-atom similarities among all compound pairs using the Tanimoto coefficient, and, in the case of MDS profiling, translates this information to coordinates on a scatter plot. To account for the relatively small library of molecules we are comparing, we employed a Tanimoto coefficient cutoff of 0.4 for determining clustering hierarchies, which might otherwise be considered too lenient for analyzing larger, more diverse datasets. The resultant MDS plot reinforced several trends seen within the tested PARPi structures (**Supplementary Fig. 7**). More condensed structures, such as 3-AB, 1,5-isoquinolinediol, and veliparib, were clustered in one quadrant, while elongated structures, such as MC3474, MC2050, and AZD5305, were enriched in another quadrant. Of note, the two reported PARP1-selective inhibitors, AZD5305 and NMS-P118, were almost superimposed, suggesting their structures are highly similar.

Our chosen similarity cutoff generated 16 similarity clusters, which successfully identified two major groupings: benzimidazole carboxamide derivatives (niraparib-like, cluster 4) and olaparib-like pthalazinone derivatives (cluster 6; **Supplementary Fig. 7**). NMS-P118, veliparib, and INO-1001 were clustered adjacent to niraparib derivatives, while AZD5305 was in between the two large clusters – indicating that its structure contains elements from both subgroups. To understand the relationship of structural similarity to binding selectivity, we then overlayed the clustering data onto our two-dimensional analysis of PARP1 and PARP2 binding (**Fig. 4**). This visualization demonstrated a clear segregation of PARPi selectivity based on their inherent chemical structures. Earlier generation PARPi with smaller pharmacophores were generally the least potent binders of PARP1 and PARP2 (clusters 9- 15). More importantly, there was a clear deviation between niraparib-like molecules and those related to olaparib. The former, which included closely related INO-1001 and NMS-P118, generally had >10-fold preference for PARP1, while the latter trended towards PARP2 selectivity and was punctuated with AZD-2461 and fluzoparib being the most PARP2-selective. Likewise, AZD5305, which bears similarity to both series, is uniquely selective for PARP1. Overall, the results strongly suggest that the two prominent series of PARPi have inherent binding biases toward either protein, which may be valid starting points for the next generation of highly selective drugs.

**Figure 4.**
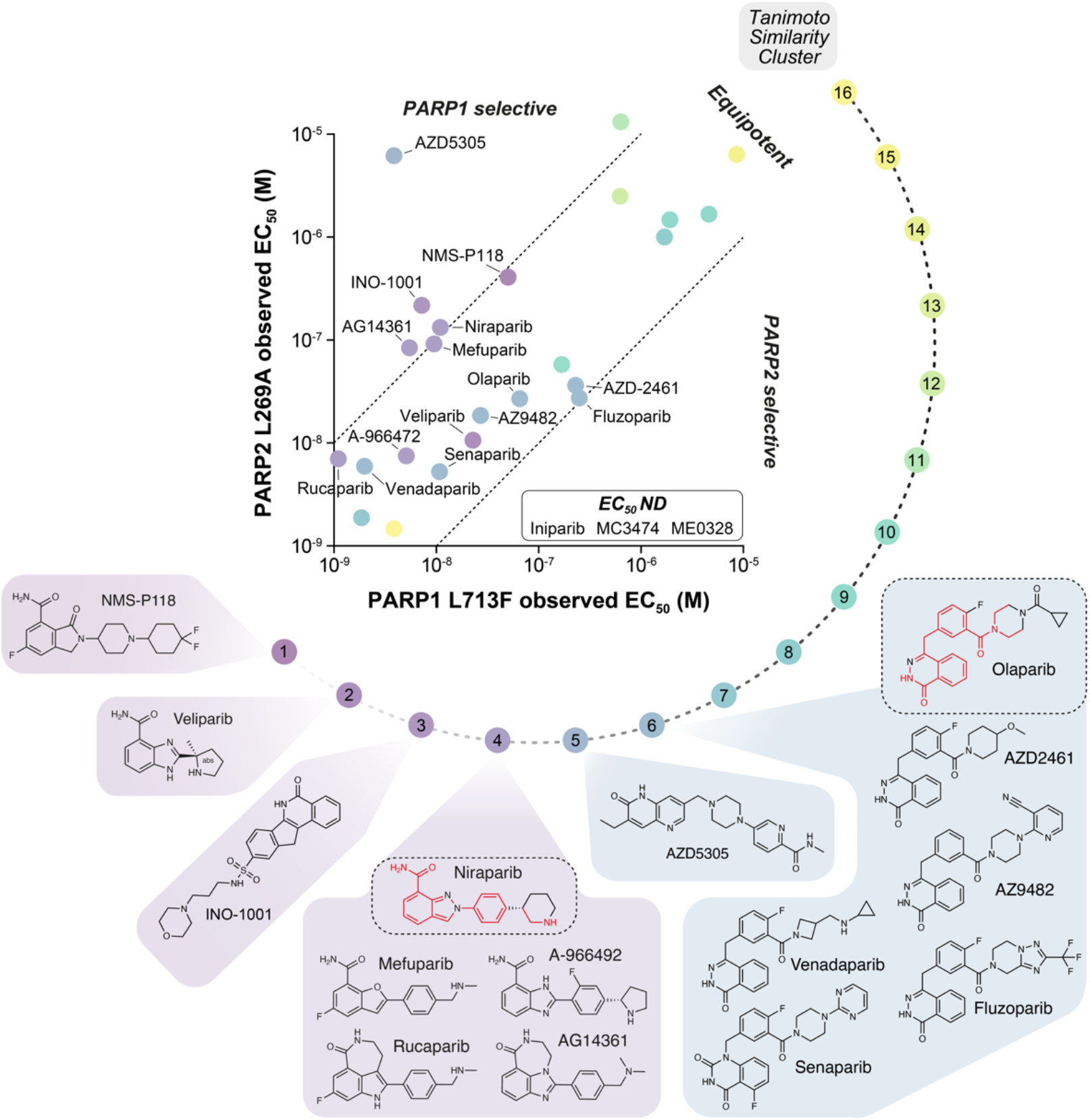
PARPi structural similarity profiling reveals pharmacophores with inherent PARP1/2 binding selectivity. Structural similarity clustering was performed with ChemMine Tools with a Tanimoto coefficient cutoff of 0.4 and overlayed onto the two-dimensional PARP1 L713F-GFP and PARP2 L269A-mCherry data (from Figure 3D). Similarity clusters (number 1-16) in color scale and PARPi structures from clusters with divergent selectivity profiles are shown. Highly similar moieties from niraparib and olaparib that are shared within relevant clusters are highlighted in red.

### HPF1 depletion modestly affects PARPi binding potency but additively promotes PARP trapping

Histone PARylation Factor 1 (HPF1) is known for its essential role in regulating PARP activity *via* redirection of PARylation activity towards histone serine residues in response to DNA damage (13). As part of this mechanism, HPF1 contributes a key glutamine residue (Glu284) to the PARP1 and PARP2 active sites that facilitates rapid PAR initiation (12, 37). Despite its low cellular abundance relative to PARP1, HPF1 effectively influences PAR synthesis by a “hit-and-run”, catalytic-like mechanism (37). Earlier *in vitro* studies have suggested that the presence of HPF1 slightly improves affinity of PARPi for PARP1 but not PARP2 (14), however its impact on PARPi binding in cells has not been comprehensively studied. To determine how HPF1 influences PARPi binding to PARP1/2, we complemented the duplexed PARP1/2 CeTEAM biosensor cells with doxycycline-regulable shRNA towards HPF1 to conditionally deplete HPF1 (**Fig. 5**). First, we generated three shRNAs targeting HPF1 and validated their knockdown efficiency at both the mRNA level by RT-qPCR (each >∼90%) and protein level by Western blot analysis (each >∼80%), demonstrating significant reductions in HPF1 expression (**Fig. 5A-C**). We then selected one hairpin (shHPF1 #1) for functional characterization.

**Figure 5.**
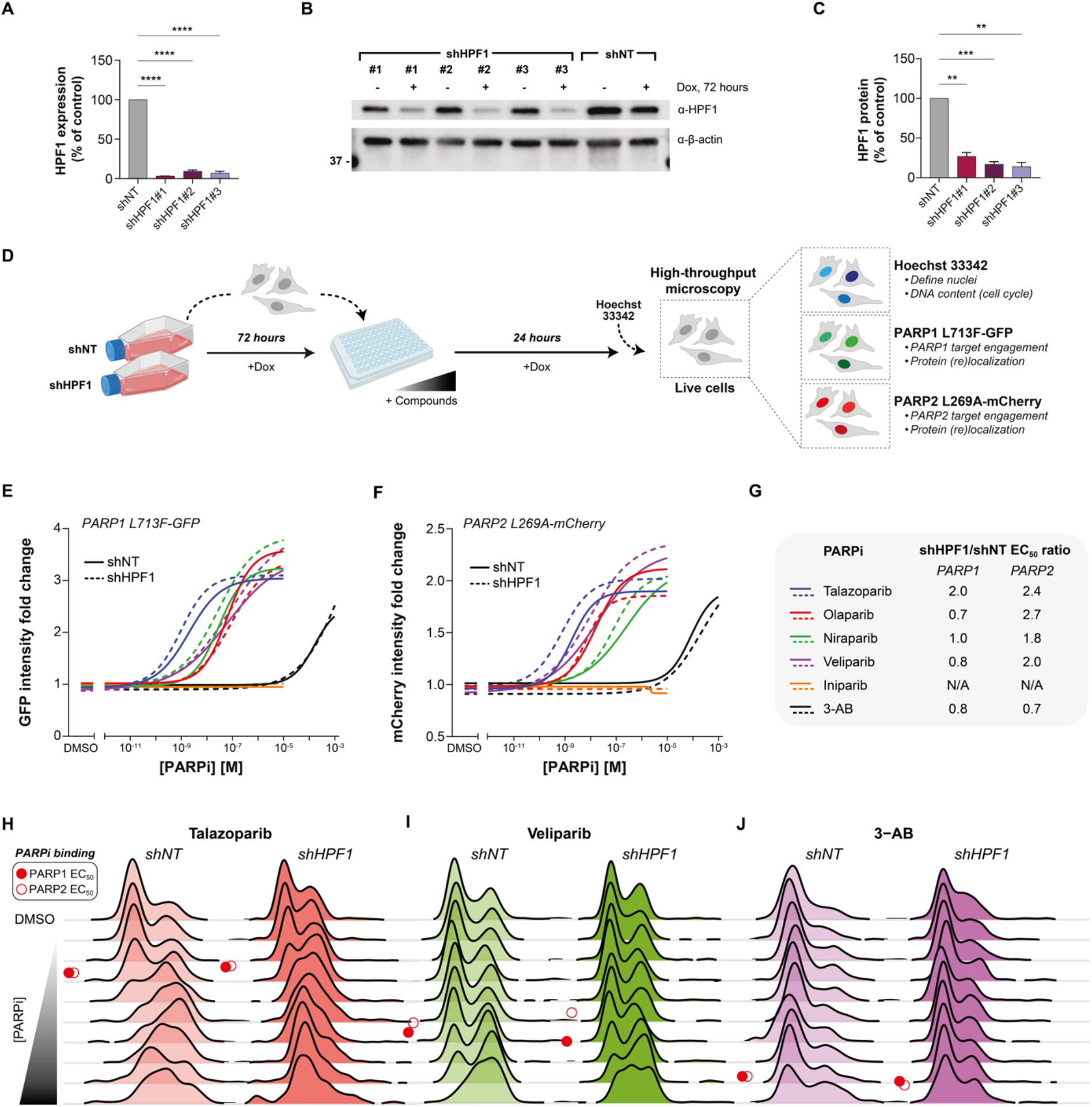
An integrated CeTEAM-genetics approach to assess the influence of HPF1 on intracellular PARPi binding and trapping. **A.** RT-qPCR analysis of gene silencing efficiency by shRNAs targeting *HPF1* (shHPF1#1, shHPF1#2, or shHPF1#3) compared to non-targeting control (shNT). Gene expression levels were normalized to *β-actin* and expressed as percentage of shNT control. Means ± SD from n=2 replicates. **** – *p* < 0.0001 as determined by one-way ANOVA with Dunnett’s post-test. **B.** Representative HPF1 protein knockdown after 72 hours DOX by Western blot. **C.** Quantification of HPF1 protein levels post-knockdown (related to **B**). Means ± SD from n=3 replicates. ** – *p* < 0.01 and *** – *p* < 0.001 as determined by one-way ANOVA and Dunnett’s post-test. **D.** Experimental set-up for live cell tracking PARP1 (GFP) and PARP2 (mCherry) target engagement and PARPi-induced replication stress (cell cycle, Hoechst) by fluorescent microscopy following HPF1 knockdown. **E.** PARP1 L713F-GFP intensity fold change in control (solid line) or HPF1 knockdown (dashed line) cells after talazoparib (blue), olaparib (red), niraparib (green), veliparib (purple), iniparib (orange), or 3-AB (black) gradients for 24 hours. Data normalized to DMSO control. Lines-of-best-fit from n=3 experiments. **F.** PARP2 L269A-mCherry intensity fold change, as in **E**. **G.** shHPF1/shNT EC_50_ value ratios (fold change) of talazoparib, olaparib, niraparib, veliparib, iniparib, and 3-AB stabilization towards PARP1 (L713F-GFP) or PARP2 (L269A-mCherry). **H.** Representative cell cycle profile dynamics of control (shNT) or shHPF1 cells after a talazoparib, veliparib (**I**), or 3-AB gradient (**J**) for 24 hours, represented as a Ridgeline plot (related to **E** and **F**). Red circles – approximate stabilization EC_50_ values for PARP1 (closed) or PARP2 (open) biosensors. N/A – not applicable

We then assessed how HPF1 knockdown affected the binding dynamics of talazoparib, olaparib, niraparib, veliparib, 3-AB, and iniparib (**Fig. 5D-G**). PARPi binding curves for a control hairpin (shNT) were similar to earlier profiles (**Fig. 1E and F**). As expected, both biosensors were unaffected by iniparib irrespective of HPF1 levels. Like data with purified proteins, HPF1 had subtle effects on observed binding of other PARPi to PARP1 and PARP2. Observed stabilization of the PARP1 biosensor was generally unchanged with clinical grade inhibitors while PARP2 binding was generally enhanced by ∼2-3-fold following HPF1 depletion (**Fig. 5G and Table 3**). Notably, talazoparib binding was improved by ∼2-fold by HPF1 depletion for both PARP1 and PARP2. Thus, HPF1 can influence PARPi binding to PARP1 and PARP2 in cells, which has important implications for intracellular PARPi selectivity. Additionally, our data demonstrates that genetic components can be layered onto CeTEAM-based assays to investigate the contribution of other proteins to observed binding potencies of a given target.

**Table 3.** Comparison of PARPi EC_50_ values in shNT or shHPF1 cells.

| **PARP1 L713F-GFP EC<sub>50</sub>** |  |  |  |  |
| --- | --- | --- | --- | --- |
| *PARPi* | *shNT (nM)* | *shHPF1 (nM)* | *logEC<sub>50</sub> p value\** | *shHPF1/shNT EC<sub>50</sub> ratio* |
| Talazoparib | 1.949 | 0.963 | 0.0209 | 2.0 |
| Olaparib | 63.80 | 30.96 | 0.4754 | 0.7 |
| Niraparib | 28.49 | 27.60 | 0.9528 | 1.0 |
| Veliparib | 63.47 | 81.47 | 0.7858 | 0.8 |
| Iniparib | NA | NA | NA | NA |
| 3-AB | 147,300 | 187,000 | 0.3837 | 0.8 |
| **PARP2 L269A-mCherry EC<sub>50</sub>** |  |  |  |  |
| *PARPi* | *shNT (nM)* | *shHPF1 (nM)* | *logEC<sub>50</sub> p value\** | *shHPF1/shNT EC<sub>50</sub> ratio* |
| Talazoparib | 1.819 | 0.7461 | 0.0180 | 2.4 |
| Olaparib | 16.01 | 5.993 | 0.0155 | 2.7 |
| Niraparib | 140.2 | 78.68 | 0.6835 | 1.8 |
| Veliparib | 18.63 | 9.159 | 0.3355 | 2.0 |
| Iniparib | NA | NA | NA | NA |
| 3-AB | 145,100 | 206,900 | 0.6325 | 0.7 |

\* – two-tailed, unpaired t test; others are not significant

In addition to assessing differences in target engagement following HPF1 knockdown, we sought to evaluate its impact on PARP inhibitor trapping effects. As HPF1 binds PARP1/2 following HD domain destabilization (12), HPF1 depletion should theoretically increase PARP trapping on DNA by increasing flexibility between the HD and catalytic domains. Indeed, as seen previously (16), HPF1 depletion induced an S/G2-phase delay consistent with PARP trapping-related replication stress (**Fig. 5H-J and Table 4**). HPF1 depletion-related S/G2 character appeared to additively compound the effects of all tested trapping PARPi, whereas profiles for iniparib and 3-AB only exhibited the basal increase from HPF1 knockdown (**Fig. 5H-J and Supplementary Fig. 8**). Overall, the data suggests that HPF1 knockdown has an additive effect on PARPi-induced trapping – presumably by already retaining a fraction of the PARP pool onto DNA.

**Table 4.** Comparison of shNT and shHPF1 trapping EC_50_ values by PARPi.

| *PARPi* | *shNT (nM)* | *shHPF1 (nM)* | *logEC<sub>50</sub> p value\** | *shHPF1/shNT EC<sub>50</sub> ratio* |
| --- | --- | --- | --- | --- |
| Talazoparib | 0.29 | 0.09 | 0.1001 | 3.2 |
| Olaparib | 14.4 | 1.89 | 0.1382 | 7.6 |
| Niraparib | 7.85 | 1.74 | 0.1295 | 4.5 |
| Veliparib | 709.1 | 55.0 | 0.0745 | 12.9 |
| Iniparib | NA | NA | NA | NA |
| 3-AB | NA | NA | NA | NA |

\* – two-tailed, unpaired t test; others are not significant

## Discussion

Poly-ADP-ribose polymerases (PARPs) are crucial players in cellular DNA repair and attractive drug targets. However, PARP1 and PARP2 have highly homologous catalytic domains, so truly selective PARPi have historically been difficult to identify. Traditional selectivity profiling is done with purified proteins, but this approach is highly dependent on the experimental conditions and omits several factors that can influence ligand binding, such as post-translational modifications (PTMs), biomolecular interaction partners, and general space constraints within the intracellular environment. Thus, cell-based selectivity assays can afford novel insights to drug interactomes and overcome some limitations of biochemical approaches (11). Here, we evaluated PARPi binding to PARP1 and PARP2 using a multiplexed cellular target engagement by accumulation of mutant (CeTEAM) approach (17) to generate live-cell selectivity and pharmacology profiles.

Overall, we found most PARPi are roughly equipotent for both PARP1 and PARP2 in cells. Among these was the next generation PARPi, senaparib, which has recently shown efficacy as a first-line option for advanced ovarian cancer (38, 39). Despite its promise in the clinic, relatively little is known about its potency and selectivity. By duplexed CeTEAM assay, we saw robust binding to both PARP1 and PARP2 (10.7 and 5.3 nM, respectively), similar to other potent, clinical-grade PARPi, thereby supporting its efficacious clinical activity.

There were also PARPi with divergent selectivity (arbitrary selectivity ratio >10-fold), but only with preference towards PARP1 – including niraparib, INO-1001, AG14361, mefuparib, NMS-P118, Tankyrase-IN-2, and AZD5305. Structural similarity analysis indicated that all these molecules, except for Tankyrase-IN-2, contain similar elements imparting preference for PARP1 binding. Most of these are extended benzimidazole carboxamide derivatives, like niraparib, or possess similarly elongated moieties, suggesting that this characteristic may be critical for disfavored binding to PARP2. Indeed, it has been reported that niraparib can sterically clash with the N-terminus of αF in the PARP2 HD domain (23), so it would not be farfetched to assume that similar steric interactions also affect these other molecules, lead to markedly better binding to PARP1 compared to PARP2.

The discrepancies in selectivity of these molecules were unexpected given the abundance of *in vitro* inhibition data supporting their equipotency. However, accurate measures of PARP1 and PARP2 inhibition are difficult to attain due to their ability to automodify themselves, resulting in a tight binding limit problem (which likely underestimates the actual potency of the inhibitor, (5)). On the other hand, measures of PARPi binding by fluorescence polarization has indicated that many niraparib-like molecules are more PARP1 selective (26), suggesting that it could be more selective of a binder than originally thought.

AZD5305 has been touted as one of the first truly PARP1-selective inhibitors to date (25, 26), and our datasets with live cell CeTEAM selectivity profiling confirm its high selectivity for PARP1 over PARP2 by a remarkable ∼1600-fold (3.9 nM *vs* 6.2 µM, respectively). Of all inhibitors tested, it was the only one that had such a high selectivity. It was recently reported that AZD5305 can still allosterically trap PARP2 onto DNA at higher concentrations (>1 µM), suggesting that it binds PARP2 above 1 µM and could influence intracellular selectivity and toxicity profiles (23). Indeed, binding and trapping of PARP2 has been implicated as the driver of hematological toxicities in patients treated with current PARPi (9, 10). However, based on our findings, there is a significant intracellular selectivity window (at least 1000-fold), implying that this can be largely avoided in a clinical setting. Thus, AZD5305 should be a promising molecule for selective PARP1 targeting and validates the use of a duplexed PARP1/2 CeTEAM assay to screen for novel, highly selective PARPi.

In addition to allosteric modulation of the HD domains within PARP1 and PARP2, trapping is believed to be primarily driven by inhibitory potency – as inhibition also blocks PARP1/2 autoPARylation, thereby promoting their retention on DNA (5, 15, 40, 41). In agreement with this notion, we generally saw that trapping correlates highly with binding potency and particularly when targeting PARP1/2 equipotently. Veliparib and, to a lesser extent, venadaparib were the clear exceptions to this trend, as they were the only potent PARPi that weren’t potent trappers. In the case of veliparib, this unique feature is likely driven by its considerably smaller size, which is unable to directly contact the HD domains and further promote PARP retention on DNA, thereby promoting its release (23, 41). Interestingly, significant trapping phenotypes by veliparib arise near the binding saturation point of the PARP1/2 biosensors, suggesting that its trapping effect is primarily driven by inhibition. In agreement with earlier reports (25), we also see that AZD5305 induces PARP trapping but to a lesser extent than most equipotent, clinical-grade PARPi (trapping EC_50_: ∼61 nM). This is conceivably due to the lack of extensive PARP2 trapping, which occurs more readily with inhibitors equipotently targeting PARP1 and PARP2. Still, the high trapping potency of AZD5305 underscores the significant contribution of higher intracellular PARP1 abundance to overall PARP trapping phenotypes.

Another surprise was that the early generation PARPi, INO-1001 (Inotek/Genentech) – which was halted after Phase 1 clinical trials (42), was a potent PARP1/2 binder with impressive selectivity for PARP1 over PARP2 (∼30-fold; 7.9 *vs* 217.2 nM, respectively). Notably, the INO-1001 selectivity profile was like that of niraparib, perhaps due to structural similarity, although insights to its PARP binding modality are unavailable for direct comparisons. Like AZD5305, INO-1001 also induced potent PARP trapping phenotypes (∼60 nM), but to a lesser extent than other clinical-grade inhibitors. Interestingly, dose-limiting toxicities of INO-1001 were hematological, which may, in retrospect, relate to inhibition or trapping of PARP2. It is unclear why clinical development was ceased, as no serious adverse effects were reported from the clinical trials and pharmacokinetics data was promising, although concerns about liver toxicity may have played a role (43, 44). Nevertheless, our data suggests it might be worth considering INO-1001 as a tool for studies differentiating PARP1 and PARP2 biology.

Besides multiplexing drug biosensors, we added to the capabilities of CeTEAM by layering on a genetic perturbation. Here, we sought to understand how HPF1, a critical component of the PARP1 and PARP2 catalytic sites, could influence binding of PARPi in cells. HPF1 was previously found to increase binding affinity of some PARPi for PARP1 but not PARP2 (5, 15). In our experiments with PARP1 L713F and PARP2 L269A biosensors, we instead saw generally improved PARPi binding when HPF1 was absent, particularly for PARP2. These discrepancies could be due to the nature of our PARP variants, as HPF1 preferentially binds PARP1/2 when the HD is in the open conformation and our mutants are constitutively flexible in the HD (12). Thus, HPF1 and ligand binding could have a compound effect on PARP biosensor stability. In line with this, as before (16), we saw that knockdown of HPF1 induced a PARP trapping-like phenotype, which could relate to how HPF1 affects PARP1/2 allostery and release from DNA. The HPF1 knockdown yielded an additive effect on PARP trapping following addition of PARPi, again suggesting that both factors contribute to HD conformational dynamics and, thus, dictate PARP1/2 retention.

In summary, we expand the capabilities of CeTEAM by both multiplexing PARP1 and PARP2 drug biosensors, while also incorporating knockdown of HPF1 to discern effects on PARPi binding and PARP trapping. The expansion into multiplexed CeTEAM biosensors opens the door for higher throughput intracellular selectivity profiling assays, yielding potentially more accurate selectivity data overall. Meanwhile, the integration of genetic perturbations into CeTEAM platforms can facilitate the discovery of novel regulators of drug-target interactions and expand the scope of drug interactomes. In the context of PARP inhibitor development, our duplexed PARP1/2 CeTEAM platform should be a useful tool to aid the discovery of next generation, PARP1- or PARP2-selective inhibitors.

## Experimental procedures

### Cell lines and culturing conditions

U-2 OS, osteosarcoma cells, HEK293T, embryonic kidney epithelial cells, and CCRF-CEM, human T lymphoblasts, were obtained from the American Type Culture Collection (ATCC, Manassass, VA, USA). U-2 OS and HEK293T cells were cultured in DMEM high glucose GlutaMAX medium (Thermo Fisher Scientific) supplemented with 1% Penicillin-Streptomycin (Thermo Fisher Scientific) and 10% fetal bovine serum – FBS (Thermo Fisher Scientific), where CCRF-CEM cells were cultured in RPMI 1640 Medium, GlutaMAX™ Supplement (Thermo Fisher Scientific) supplemented with 1% Penicillin-Streptomycin (Thermo Fisher Scientific) and 10% fetal bovine serum. For *in vitro* fluorescent microscopy read-outs, DMEM, high glucose, no glutamine, no phenol red media (Thermo Fisher Scientific) supplemented with 1x GlutaMAX, 1% Penicillin-Streptomycin (Thermo Fisher Scientific) and 10% fetal bovine serum – FBS (Thermo Fisher Scientific) was used. Cell cultures were maintained at 37°C with 5% CO2 in a humidified incubator.

### Antibodies and chemicals

anti-mCherry (rabbit polyclonal, cat. No. PA5-34974), donkey anti-rabbit Alexa Fluor 568 (cat. No. A10042) and goat anti-rabbit Alexa Fluor 488 (cat. No. A11008) were obtained from Thermo Fisher, anti-PARP2 (rabbit polyclonal, cat. No. 55149-1-AP) and anti-PARP1 recombinant antibody (rabbit recombinant, cat. No 80174-1-RR-20) were acquired from Proteintech, anti-GAPDH (polyclonal rabbit, cat. No. ab9485) was purchased by Abcam, anti β-actin (mouse monoclonal, A5441) was obtained from Sigma, anti-HPF1 (rabbit monoclonal, 90876), was acquired from Cell Signaling, anti-PARP1 (mouse monoclonal, sc8007) and anti-GFP (mouse monoclonal, sc-9996) were purchased from Santa Cruz Biotechnologies. Doxycycline hydrochloride (Sigma-Aldrich) was dissolved in MilliQ water (2 mg/mL) and used at 0.75 μg/mL. Talazoparib, niraparib, olaparib, veliparib, AZ9482, senaparib, tankyrase-IN2 and INO-1001 (MedChemExpress) were dissolved in DMSO to a stock of 10 mM. Iniparib (MedChemExpress) was dissolved in DMSO to a stock of 20 mM. 3-aminobenzamide (3-AB; Sigma-Aldrich) was dissolved in DMSO to a stock of 100 mM. A-966492, pamiparib, PJ34, MC2050, MC3474, mefuparib, AZD5305, and rucaparib were kindly provided by the Rotili Lab and dissolved in DMSO at 10 mM. NMS-P118, venadaparib, BYK204165, AG14361, E7449, AZD-2461, fluzoparib, and 1,5-isoquinolinediol were provided as 10 mM stocks in DMSO from the SciLifeLab Compound Center (refer to **Table S2**). Bisbenzimide H 33342 trihydrochloride (Hoechst 33342, Thermo Fisher) was dissolved in MilliQ water to a stock of 20 mg/mL.

### Recombinant DNA cloning

PARP2 L269A was subcloned (**Table S1**) from pENTR1a-PARP2 L269A-GFP (6, 17) into pENTR1a-C-mCherry by flanking SalI/NotI restriction sites and subsequently transferred to pLenti CMV Hygromycin DEST (Addgene plasmid #17454) or Ef1a-Tta3G-P2A-Blast (45). pENTR1a-PARP1 L713F-C-GFP was transferred to pLenti CMV Blast DEST (Addgene plasmid #17451).

Short hairpin RNA (shRNA) HPF1#1, #2 and #3, and NT (non-targeting) were oligo annealed into a prSITEP-TetR-Neomycin-akaLuc. The shRNA system backbone of the pRSITEP-U6Tet-(sh)- EF1-TetRep-2A-Puro-P2A-RFP670 plasmid (46) was modified to include a Tet-Repressor for controlled shRNA expression via doxycycline, a Neomycin resistance gene for antibiotic selection, and a luminescent reporter, akaLuc. The entire sequence is transferable using XbaI/SalI restriction sites. All subcloning into entry vectors was validated by automated sequencing; while shuttling into destination vectors was performed with Gateway LR Clonase II (ThermoFisher Scientific) and positive clones were confirmed by colony PCR. All plasmids, synthetic genes, and primers (Eurofins Genomics) are listed in **Table S1**.

### Lentivirus production and transduction

Lentiviral production was performed following transfection of third generation lentiviral packing vectors by calcium phosphate precipitation. prSITEP Neo, pLenti CMV Hygro, pLenti CMV Blast, or pLenti CMV Puro lentiviral constructs were co-transfected with lentiviral packaging vectors (Gag-Pol, Rev, and VSV-G envelope) into subconfluent HEK293T cells. Viral particles were harvested at 48- and 72-hours post-transfection, and target cells were transduced at 1:1 dilution of lentivirus and fresh, complete medium in the presence of polybrene (8 μg/mL). Forty-eight hours post-transduction, target cells were re-plated at low density in the presence of G418/neomycin (Sigma-Aldrich, 450 μg/mL for six days; pINDUCER20 – Addgene plasmid #44012), puromycin (Sigma-Aldrich, 1 μg/mL for three days; pLenti CMV Puro – Addgene plasmid #17452), blasticidin (Sigma-Aldrich, 5 μg/mL for four days; pLenti CMV Blast – Addgene plasmid #17451), or hygromycin (Sigma-Aldrich, 100 µg/mL for five days; pLenti CMV Hygro – Addgene plasmid #17454) that was replenished at three-day intervals.

### Reverse transcription quantitative PCR (RT-qPCR)

U-2 OS cells were plated at 60,000 cells/well in 6-well plates in the absence or presence of 0.75 μg/mL DOX. After 72 hours, the cells were harvested with TRIzol (Thermo Fisher Scientific). RNA was purified with the Direct-zol RNA MiniPrep kit (Zymo Research) according to the manufacturer’s instructions and quantified on a De Novix DS-11 FX Spectophotometer/Fluorometer. cDNA was then generated with the iScript cDNA Synthesis Kit (Bio-Rad) according to the manufacturer’s instructions. qPCR was performed with 2.5 ng cDNA per sample and iTaq Universal SYBR Green Supermix (Bio-Rad) using a Bio-Rad CFX96 Real-Time PCR Detection System. Relative quantity of target genes was calculated using the ΔΔCt method *via* normalization to *β-actin*. All qPCR primers are listed in **Table S1** and were ordered from Eurofins.

### Western blotting

Cells were collected by trypsinization and lysed in RIPA buffer (20 minutes on ice with occasional mixing), and the clarified lysate was supplemented with Laemmli buffer (1x final). Following heating for 5 minutes at 95°C, the samples were either directly loaded for electrophoresis or frozen at -20°C for later use. Protein samples were separated on 4-20% gradient Mini-PROTEAN gels (Bio-Rad) or Midi-PROTEAN gels (Bio-Rad) prior to transferring onto 0.2 μm nitrocellulose with a Trans-Blot Turbo Transfer System (Bio-Rad). After blocking with ROTI®Block (Carl Roth) for 1 hour at room temperature primary antibodies were applied in 1x ROTI®Stock TBST (Carl Roth) at specified concentrations and left to incubate on a shaker at 4°C overnight. The primary antibodies used included anti-GFP probe (mouse monoclonal, 1:500), anti-mCherry (rabbit polyclonal, 1:2000), anti-PARP1 (mouse monoclonal, 1:500), anti-PARP2 (rabbit polyclonal, 1:2000), anti-GAPDH (rabbit, 1:2500), anti-β-actin (mouse monoclonal, 1:5000), and anti-HPF1 (rabbit, 1:1000). LI-COR secondary antibodies were diluted in 1x ROTI®Stock TBST (Carl Roth) at 1:10,000 and incubated at room temperature for one hour. The blots were then imaged using a LI-COR Odyssey Fc system and analysed with Image Studio Software (LI-COR).

### Live cell fluorescence microscopy

For PARP L713F and PARP2 L269A experiments in live U-2 OS cells, 44,000 cells were plated in transparent, clear bottom 96-well plates (BD Falcon) on day 0 in complete medium. The following day medium was changed to complete DMEM phenol-free medium, and inhibitors were added to their indicated final concentrations in complete DMEM phenol-free medium (final DMSO 0.1% or 1% [3-AB] v/v). After 16 or 24 hours, cell-permeable Hoechst 33342 was added to a final concentration of 1μg/mL for 30 minutes prior to imaging.

For shHPF1 experiments in live U-2 OS cells, Dox 0.75ug/mL was added 72h prior the start of the experiment. After this period, cells were trypsinized and 44,000 cells were plated in transparent, clear bottom 96-well plates (BD Falcon) on day 0 in DMEM phenol-free medium. The next day, inhibitors were added to their indicated final concentrations in complete DMEM phenol-free medium (final DMSO 0.1% or 1% [3-AB] v/v). After 24 hours, cell-permeable Hoechst 33342 was added to a final concentration of 1μg/mL for 30 minutes prior to imaging.

Imaging was performed using the CellCyteX microscope (Cytena) at 10x magnification. The microscope was maintained at 37°C with 5% CO2 in a humidified incubator. Image analysis was then performed with CellProfiler software (Broad Institute) where the mean nuclei intensity upper quartile intensity mask of both GFP and mCherry were used to understand the fold changes and the integrated Hoechst intensity was used for cell cycle analysis.

### Small molecule screen details

#### Composition, storage, and plating of screened compounds

Hit molecules from an earlier PARP1 L713F stability screen (17) were handled and plated as before but into clear plates 96-well plates cell culture plates (Starstedt) using an Echo 550™ acoustic liquid handler (LabCyte). An overview of the small molecules screened is detailed in **Table S2**.

#### Screen execution

U-2 OS PARP1 L713F-GFP/PARP2 L269A-mCherry cells were plated at 10,000 cells per well into drug-containing assay plates (final concentration 10 µM) using a Multidrop Combi liquid dispenser (Thermo Scientific). Cells were then incubated with drugs for 16 hours at 37 °C with 5% CO_2_ in a humidified incubator. To minimize edge effects, the plates were placed in self-made humidity chambers that limited evaporation in the outer ring of wells. Imaging was performed using the CellCyteX microscope (Cytena) at 10x magnification. The microscope was maintained at 37°C with 5% CO2 in a humidified incubator. Image analysis was then performed with CellProfiler software (Broad Institute) where the mean nuclei intensity upper quartile intensity mask of both GFP and mCherry were used to understand the fold changes. The same strategy was applied to measure the dose-curves of all the non-PARPi (DC-05, A-395, ACY-1083, droxinostat, KDM5-C70, bisindolylmaleimide I (GF109203X), AZ960, Go 6983, linifanib (ABT-869), paclitaxel, PF-3758309, AT9283, azacitidine and decitabine).

### Cellular thermal shift assay (CETSA)

CCRF-CEM cells (1×10^6^ per treatment) were harvested by centrifugation at 500 x g for 5 minutes and washed twice with phosphate-buffered saline (PBS). Post-washing, cells were resuspended in 4mL Tris-buffered saline (TBS) containing a cocktail of proteasome inhibitors (EDTA-free, Roche). Cell lysis was achieved by alternately exposing cells to dry ice and absolute ethanol (VWR) for 3 minutes and to a temperature of 37°C for 3 minutes, repeating this cycle three times. Subsequently, lysates were centrifuged at 20,000 x g for 20 minutes at 4°C. The supernatant was collected and aliquoted into 100 µL fractions. Each aliquot was treated with either DMSO or 10 µM olaparib and incubated at room temperature for 20 minutes. Thermal denaturation was conducted at a range of temperatures: 37°C, 40°C, 43°C, 46°C, 49°C, 52°C, 55°C, 58°C, 61°C, 64°C, 67°C, and 70°C, each for 3 minutes. Samples were then cooled down at room temperature for 3 minutes before being centrifuged at 20,000 x g for 20 minutes at 4°C. Samples were prepared by adding Laemmli 4X sample buffer and heated to 95°C for complete denaturation. Proteins were then resolved by SDS-PAGE and transferred onto membranes for Western blot analysis, as above.

### Immunofluorescence (IF) microscopy

U-2 OS cells were seeded at a density of 4.44×10^4^ cells per well in a PhenoPlate 96-well plate (Perkin Elmer). The next day, DMSO (0.1% v/v, final) or 10 µM bisindolylmaleimide I (GF109203X) was added to each well. After 24 hours the cells were fixed with 4% paraformaldehyde (PFA - Histolab) for 15 minutes and then permeabilized with 0.1% Triton X-100 (Sigma) in PBS for 15 minutes. Afterwards, the cells were blocked with 2% bovine serum albumin (BSA - Sigma) in PBS for one hour at room temperature. The cells were then incubated with a primary anti-PARP1 recombinant antibody (rabbit recombinant), diluted 1:200 in 0.1% BSA in PBS, for two hours at room temperature. Alexa Fluor 568 and Alexa Fluor 488-conjugated secondary antibodies (anti-rabbit) were diluted 1:500 in 0.1% BSA in PBS and applied to the cells at room temperature in the dark. After incubation with secondary antibodies. Cells were stained with Hoechst at a 1:2000 dilution for 5 minutes at room temperature. Cells were imaged using a A1R+ Nikon confocal microscope system at 20x magnification. The 568-channel spectral free band (VF) ranged between 590-620 nm and the 488-channel spectral free band (VF) ranged between 490-530 nm.

### Statistical analysis

All graphing and statistical analyses were performed using GraphPad Prism V10 or R v4.1.1. Saturation curve fitting was performed using the [agonist] vs response four parameter variable slope model in GraphPad Prism. Specific post-hoc tests, variations, and statistical significances for relevant experiments are described within individual figure legends. For **Tables 3 and 4** the logEC_50_ p values were measured using unpaired, two-tailed t test assuming both populations have the same SD (standard deviation).

## Data and materials availability

All data is available in the main text and in supporting information. Materials are available from the corresponding authors (Nicholas Valerie –; Mikael Altun –) upon reasonable request.

## Supporting information

This article contains supporting information.

## Acknowledgments

We would like to express our gratitude to the SciLifeLab Compound Center, part of the Chemical Biology Consortium Sweden (CBCS), for assistance with compound management and spotting.

## Author contributions

Conceptualization (NCKV, MA), Methodology (MJP, NCKV, SA, MA), Software (AL), Validation (MJP, MA, NCKV), Formal analysis (MJP, AL), Investigation (MJP, NCKV), Resources (NCKV, MA, EF, DR), Writing – original draft (MJP), Writing – review & editing: (MJP, AL, SA, EF, DR, MA, NCKV), Visualization: (MJP, AL, NCKV, MA), Supervision (NCKV, MA, DR), Project administration (NCKV, MA), Funding acquisition (NCKV, MA, DR).

