## Supplementary Information for "Duplexed CeTEAM drug biosensors reveal determinants of PARP inhibitor selectivity"

**Table of Contents**

Supplementary tables ........................................ S3

Table S1. Oligo annealing primers, plasmids and qPCR primers ........................................ S3

Table S2. Screening compound details from the SciLifeLab Compound Center. ........................................ S4

Supplementary figures ........................................ S5

Supplementary Figure 1. Olaparib binding to PARP1 and PARP2 by cellular thermal shift assay (CETSA) ........................................ S5

Supplementary Figure 2. Dose-dependency of non-PARPi stabilization of PARP1 and PARP2 drug biosensors. A. ........................................ S6

Supplementary Figure 3. Bisindolylmaleimide I and Go 6983 are autofluorescent assay artefacts. .. S7

Supplementary Figure 4. Full dose-response curves for PARPi stabilization of PARP1 L713F-GFP and PARP2 L269A-mCherry in dual biosensor cells. ........................................ S8

Supplementary Figure 5. Evaluation of various PARPi on PARP1 L713F-GFP and PARP2 L269A-mCherry protein levels by western blot. ........................................ S9

Supplementary Figure 6. Cell cycle profile dynamics of tested PARPi. ........................................ S10

Supplementary Figure 7. 2D multidimensional scaling (MDS) of PARPi structures. ........................................ S11

Supplementary Figure 8. Influence of HPF1 depletion on PARPi cell cycle dynamics. ........................................ S12

**Supplementary tables**

**Table S1.** Oligo annealing primers, plasmids and qPCR primers

| **Oligo annealing primers** | *Sequence (5'→ 3')* |
|---|---|
| shHPF1#1 F | CCGGGCTTGGTTGTTCCAGTAGATACTCGAGTATCTACTGGAACAACCAAGCTTTTTG |
| shHPF1#1 R | AATTCAAAAAGCTTGGTTGTTCCAGTAGATACTCGAGTATCTACTGGAACAACCAAGC |
| shHPF1#2 F | CCGGGTGAAGAACTTGATCCTGAAACTCGAGTTTCAGGATCAAGTTCTTCACTTTTTG |
| shHPF1#2 R | AATTCAAAAAGTGAAGAACTTGATCCTGAAACTCGAGTTTCAGGATCAAGTTCTTCAC |
| shHPF1#3 F | CCGGGCAAGTGATGAGGAGAGACTACTCGAGTAGTCTCTCCTCATCACTTGCTTTTTG |
| shHPF1#3 R | AATTCAAAAAGCAAGTGATGAGGAGAGACTACTCGAGTAGTCTCTCCTCATCACTTGC |
| **Plasmids** | *Source* |
| pENTR1a-PARP1 L713F-GFP | Valerie., *et al.*[1] |
| pENTR1a-PARP1 WT-GFP | Valerie., *et al.*[1] |
| pENTR4-mCherry-PARP1 WT | This work |
| pLenti CMV Blast_PARP1 L713F-GFP | This work |
| pINDUCER20-PARP1 L713F-GFP | Valerie., *et al.*[1] |
| pINDUCER20-PARP1 WT-GFP | Valerie., *et al.*[1] |
| Ef1a-Tta3G-P2A-Blast_PARP1 WT_mCherry | This work |
| pLenti CMV Puro_PARP1 WT-mCherry | This work |
| pET28-PARP2 L269A | Langelier, M.-F., *et al.*[2] |
| TOPO-PARP2 L269A | Valerie., *et al.*[1] |
| pET28-PARP2 WT | Langelier, M.-F., *et al.*[2] |
| TOPO-PARP2 WT | This work |
| pENTR1a-PARP2 L269A-mCherry | This work |
| pINDUCER20-PARP2 WT-GFP | This work |
| pLenti CMV Hygro_PARP2 WT-mCherry | This work |
| pLenti CMV Hygro_PARP2 L269A-mCherry | This work |
| prSITEP_Puro_akaLuc-shHPF1#1 | This work |
| prSITEP_Puro_akaLuc-shHPF1#2 | This work |
| prSITEP_Puro_akaLuc-shHPF1#3 | This work |
| prSITEP_Puro_akaLuc-shNT | This work |
| **qPCR primers** | *Sequence (5'→ 3')* |
| HPF1 F | AGAAAGTTGTGACAAAGACC |
| HPF1 R | CATCATTTCCTGAATGGGAG |
| β-actin F | CCTGGCACCCAGCACAAT |
| β-actin R | GGGCCGGACTCGTCATACT |

**Table S2.** Screening compound details from the SciLifeLab Compound Center.

| Compound | Compound ID | Batch number | Stock (mM) | Volume (nL) |
|---|---|---|---|---|
| Linifanib (ABT-869) | CBK169041 | BJ1886001 | 10 | 100 |
| Olaparib (AZD2281) | CBK277996 | BJ1886025 | 10 | 100 |
| Paclitaxel | CBK011621 | BJ1886057 | 10 | 100 |
| PF-3758309 | CBK293899 | BJ1886257 | 10 | 100 |
| Veliparib (ABT-888) | CBK277926 | BJ1886002 | 10 | 100 |
| AT9283 | CBK277923 | BJ1886053 | 10 | 100 |
| Niraparib (R-enantiomer) | CBK278031 | BJ1886189 | 10 | 100 |
| Rucaparib (phosphate) | CBK277950G | BJ1886038 | 10 | 100 |
| 5-Azacytidine | CBK041875 | BJ1886134 | 10 | 100 |
| Decitabine | CBK201329 | BJ1886078 | 10 | 100 |
| Tankyrase-IN-2 | CBK506899 | DO8144443 | 10 | 100 |
| Decitabine | CBK201329 | DO8144447 | 10 | 100 |
| BYK204165 | CBK290351 | DO8144458 | 10 | 100 |
| Olaparib (AZD2281) | CBK277996 | DO8144485 | 10 | 100 |
| AG14361 | CBK290655 | DO8144498 | 10 | 100 |
| Bisindolylmaleimide I | CBK040989 | DO8144499 | 10 | 100 |
| E7449 | CBK308723 | DO8144504 | 10 | 100 |
| AZ960 | CBK288327 | DO8144511 | 10 | 100 |
| Go 6983 | CBK290912 | DO8144531 | 10 | 100 |
| INO-1001 | CBK506935 | DO8144542 | 10 | 100 |
| PJ34 (hydrochloride) | CBK308811C | DO8144559 | 10 | 100 |
| AZD-2461 | CBK290993 | DO8144576 | 10 | 100 |
| Rucaparib | CBK277950 | DO8144602 | 10 | 100 |
| Niraparib (tosylate) | CBK278031G | DO8144605 | 10 | 100 |
| Rucaparib (Camsylate) | CBK277950H | DO8144641 | 10 | 100 |
| Fluzoparib | CBK506988 | DO8144643 | 10 | 100 |
| 1,5-Isoquinolinediol | CBK507027 | DO8144711 | 10 | 100 |
| Talazoparib | CBK309483 | DO8144726 | 10 | 100 |
| Pamiparib | CBK506999 | DO8144660 | 10 | 100 |
| A-966492 | CBK507022 | DO8144701 | 10 | 100 |
| Senaparib | CBK506721 | DO8144123 | 10 | 100 |
| Veliparib (dihydrochloride) | CBK277926C | DO8144731 | 10 | 100 |
| ACY-1083 | CBK506771 | DO8144206 | 10 | 100 |
| A-395 | CBK506730 | DO8144137 | 10 | 100 |
| Veliparib (ABT-888) | CBK277926 | DO8144161 | 10 | 100 |
| AZ9482 | CBK506867 | DO8144391 | 10 | 100 |
| Niraparib (R-enantiomer) | CBK290547 | DO8144329 | 10 | 100 |
| KDM5-C70 | CBK506855 | DO8144369 | 10 | 100 |
| Rucaparib (phosphate) | CBK277950G | DO8144062 | 10 | 100 |
| Mefuparib (hydrochloride) | CBK506689C | DO8144061 | 10 | 100 |
| Niraparib (R-enantiomer) | CBK278031 | DO8144020 | 10 | 100 |
| NMS-P118 | CBK506686 | DO8144053 | 10 | 100 |
| Niraprib (tosylate) | CBK278031 | DO8144101 | 10 | 100 |
| Venadaparib | CBK506696 | DO8144071 | 10 | 100 |
| Niraparib (hydrochloride) | CBK278031C | DO8144089 | 10 | 100 |
| Talazoparib tosylate | CBK309483G | DO8144102 | 10 | 100 |
| AZD5305 | CBK506718 | DO8144116 | 10 | 100 |

**Supplementary figures**

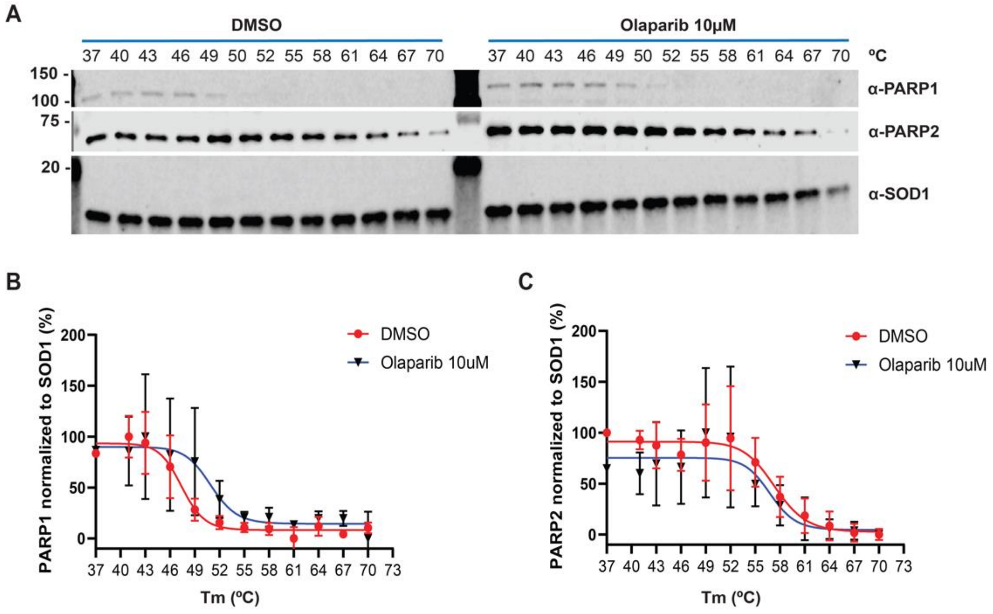


**Supplementary Figure 1. Olaparib binding to PARP1 and PARP2 by cellular thermal shift assay (CETSA). A.** Western blot analysis was employed to evaluate the thermal stability of PARP1 and PARP2 after CETSA was performed across a range of temperatures from 37ºC to 70ºC in 3ºC increments. Proteins were detected using specific antibodies against PARP1 and PARP2, with SOD1 serving as the loading control to ensure consistent protein quantification across samples. **B.** Melting profiles of PARP1 after DMSO (red) or 10 μM olaparib treatment (black/blue). Protein levels are normalized to SOD1 and set relative to 37ºC abundance (percent). **C.** Melting profiles of PARP2, as in **B**. Means from n=3 replicates ± SD with lines-of-best-fit shown in both instances.

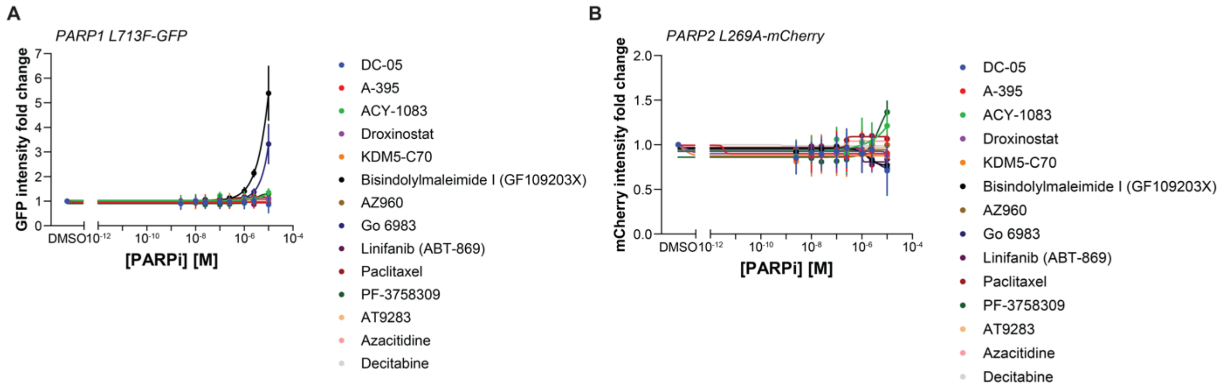


**Supplementary Figure 2. Dose-dependency of non-PARPi stabilization of PARP1 and PARP2 drug biosensors. A**. PARP1 L713F-GFP intensity fold change after non-PARPi drug gradients for 16 hours. Data normalized to DMSO control. Means from n=3 replicates ± SD with lines-of-best-fit shown. **B.** PARP2 L269A-mCherry intensity fold change, as in **A**.

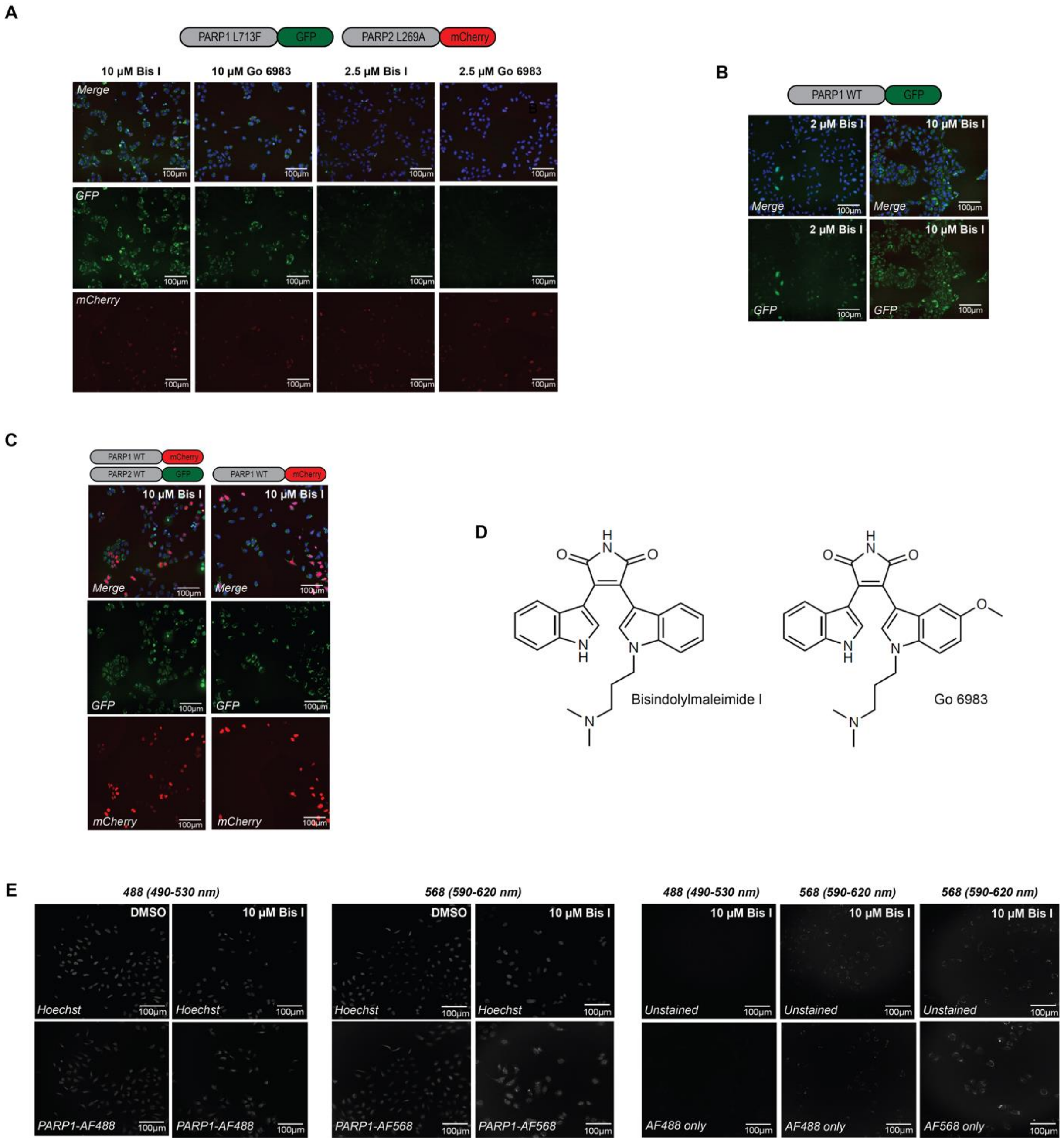


**Supplementary Figure 3. Bisindolylmaleimide I and Go 6983 are autofluorescent assay artefacts.** **A.** Live cell fluorescent microscopy of Bisindolylmaleimide I (Bis I) and Go 6983 effects on subcellular localization of PARP1 L713F-GFP and PARP2 L269A-mCherry at 2.5 or 10 µM. Distinct perinuclear accumulation of L713F-GFP is seen but not for L269A-mCherry. Scale bars = 100 µm. **B.** Live cell fluorescent microscopy of Bis I and Go 6983 effects on subcellular localization of PARP1 WT-GFP. Scale bars = 100 µm. **C.** Live-cell fluorescent microscopy of 10 µM Bis I effects on distribution of PARP1 WT-mCherry/PARP2 WT-GFP (left) or PARP1 WT-mCherry alone (right). Scale bars = 100 µm. **D.** Structures of Bisindolylmaleimide I and Go 6983. **E.** Immunofluorescence confocal microscopy assessing endogenous PARP1 localization in U-2 OS cells treated with DMSO or Bisindolylmaleimide I, using an anti-PARP1 primary antibody and Alexa Fluor 488 or Alexa Fluor 568 as secondary antibodies. Endogenous PARP1 was detected in spectrally distinct 488 (490-530 nm) or 568 (590-620) filters. Unstained cells or those with secondary antibody only were also treated with Bis I for reference. Scale bars = 100 µm.

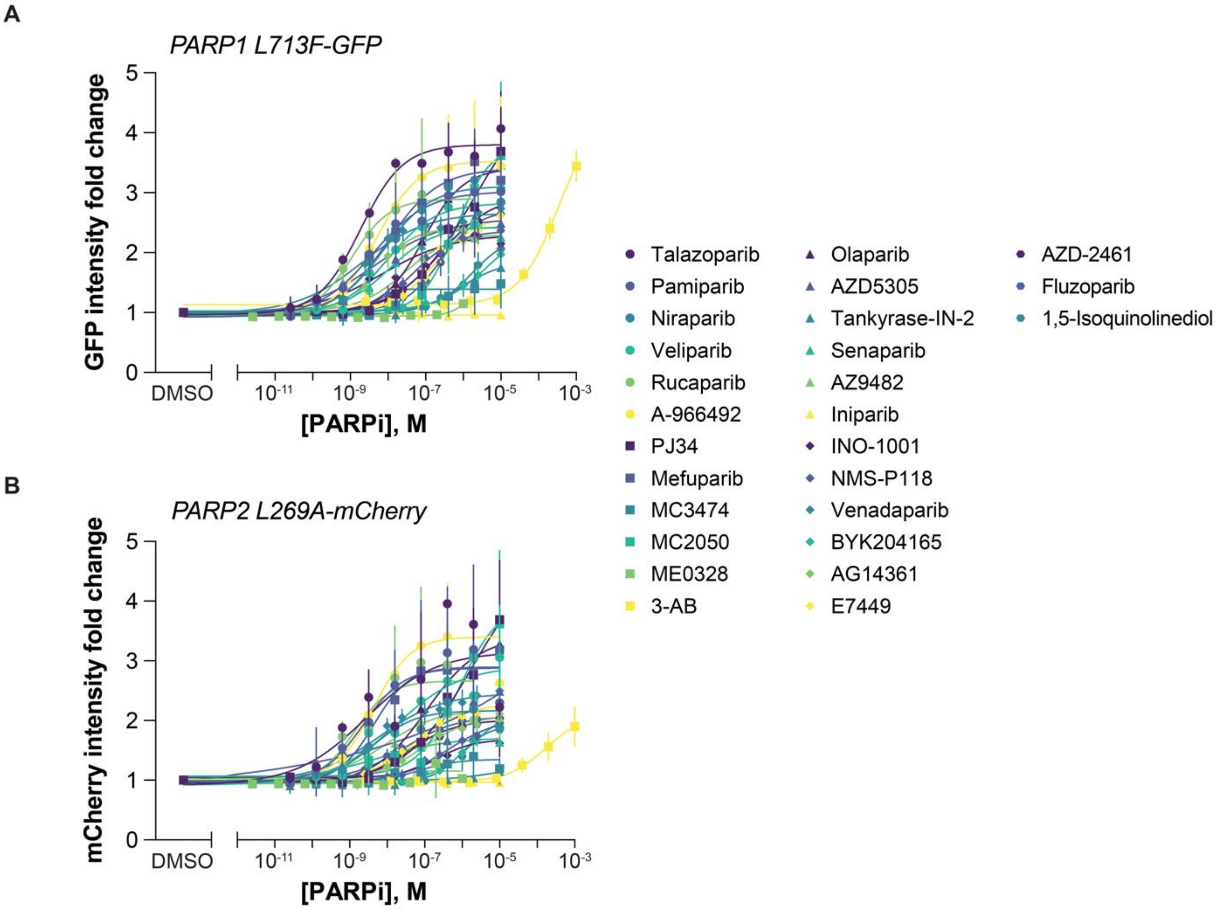


**Supplementary Figure 4. Full dose-response curves for PARPi stabilization of PARP1 L713F-GFP and PARP2 L269A-mCherry in dual biosensor cells. A.** Live-cell L713F-GFP fold change by fluorescence microscopy following a concentration gradient with the indicated PARPi for 24 hours and normalization to DMSO controls. Means ± SD (n=3) with lines-of-best-fit shown. **B.** Live-cell L269A-mCherry fold change by fluorescence microscopy, as in **A**. Means ± SD (n=3) with lines-of-best-fit shown.

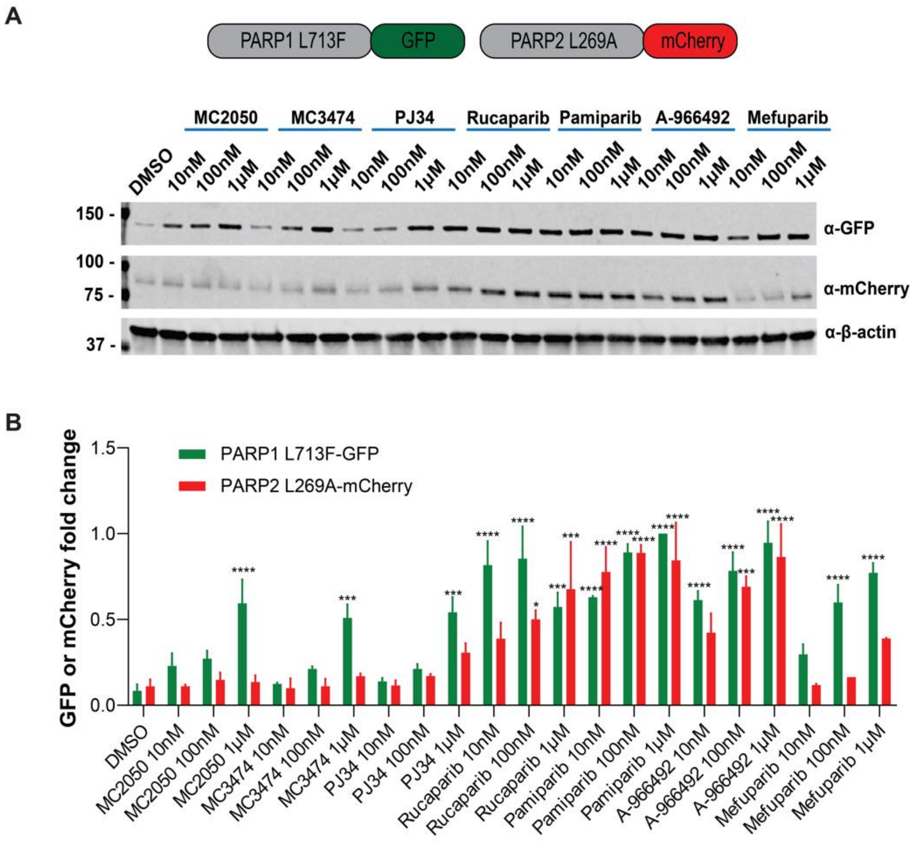


**Supplementary Figure 5. Evaluation of various PARPi on PARP1 L713F-GFP and PARP2 L269A-mCherry protein levels by western blot. A.** A representative western blot of PARP1 L713F-GFP and PARP2 L269A-mCherry abundance 24 hours after MC2050, MC3474, PJ34, rucaparib, pamiparib, A-966492, or mefuparib treatment at the indicated concentrations. **B.** Densitometric quantification of PARP1 L713F-GFP (green) and PARP2 L269A-mCherry (red) protein levels relative to β-actin and normalized to the highest value recorded within the experiment to ensure comparability across treatments. Means ± SD from n=2 replicates. Statistical significance indicated as * – $p < 0.05$, *** – $p < 0.001$, and **** – $p < 0.0001$ as determined by one-way ANOVA with Dunnett's post-test, comparing each treatment to the DMSO control (all others not significant).

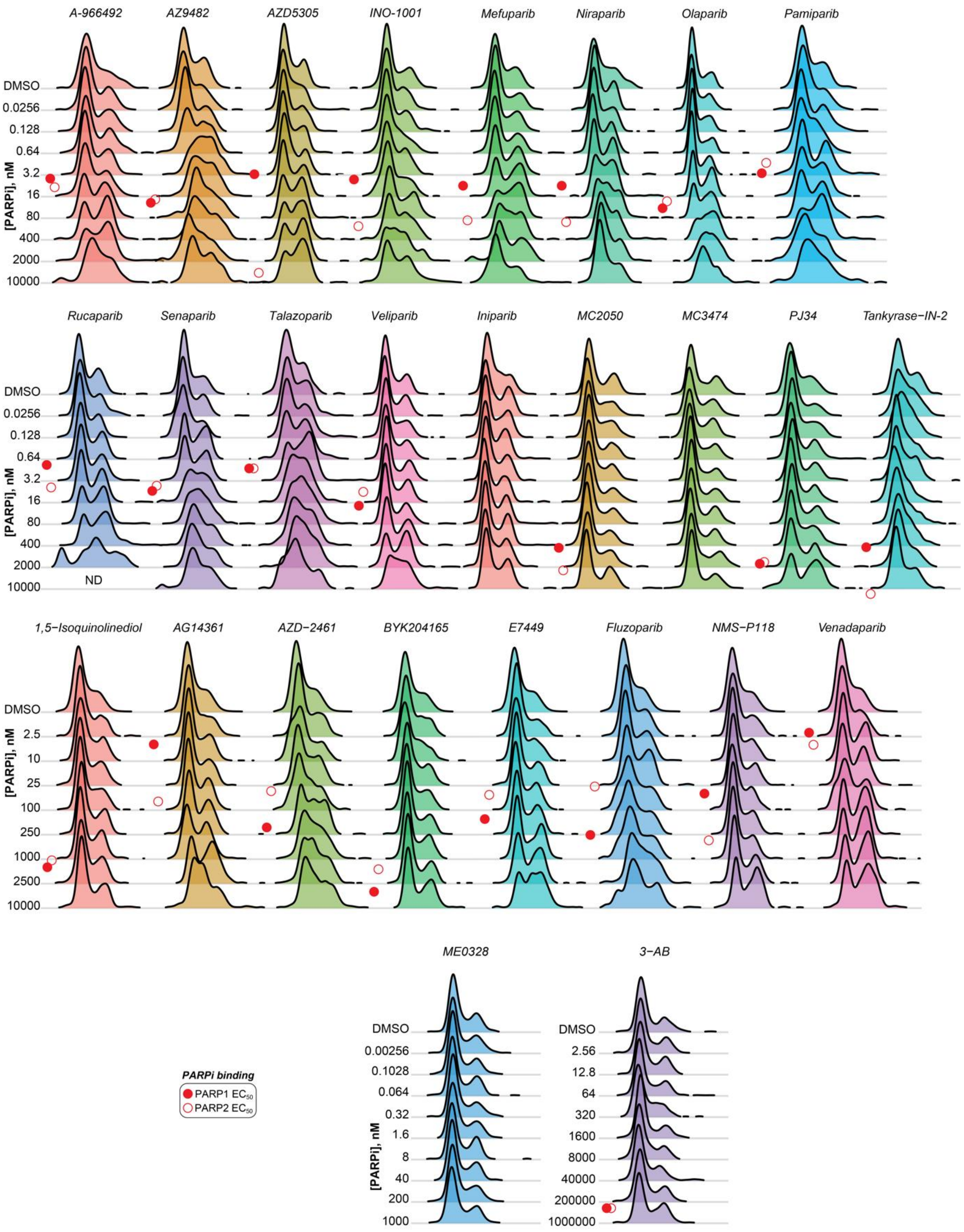


**Supplementary Figure 6. Cell cycle profile dynamics of tested PARPi.** Representative cell cycle profile dynamics from Hoechst intensity for PARPi tested by dose-response, as shown by Ridgeline plot. Concentrations of PARPi are indicated on the lefthand side of the plots. Red circles – approximate stabilization $EC_{50}$ values for PARP1 (closed) or PARP2 (open) biosensors.

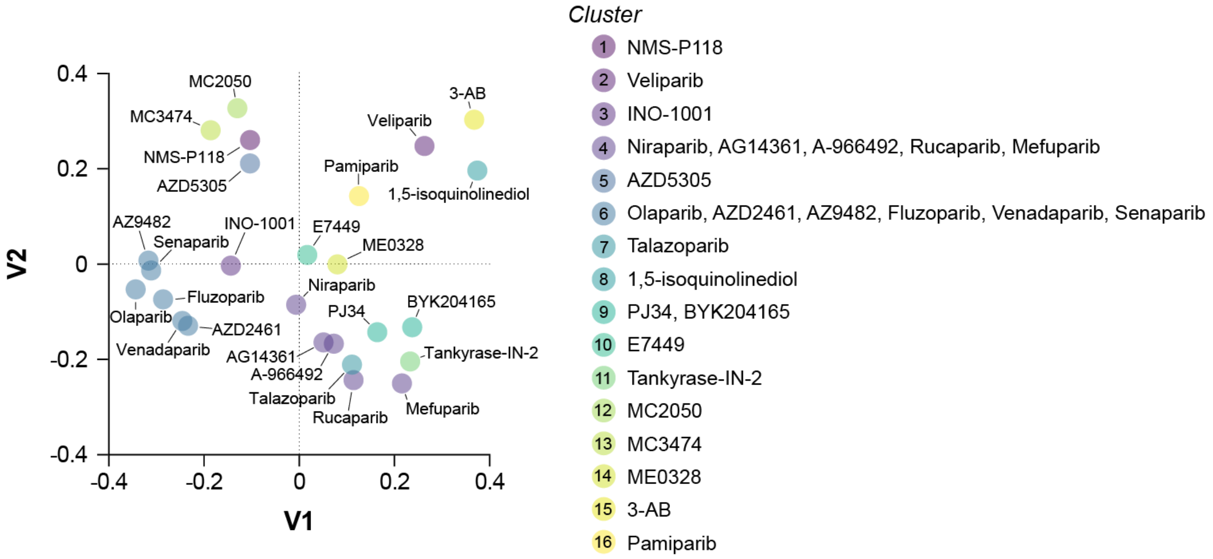


**Supplementary Figure 7. 2D multidimensional scaling (MDS) of PARPi structures.** Structural similarity clustering and MDS plotting was performed with the ChemMine Tools online server with a Tanimoto coefficient cutoff of 0.4 for determining clusters. Clusters are defined by number and associated color label with PARPi falling into each cluster labelled.

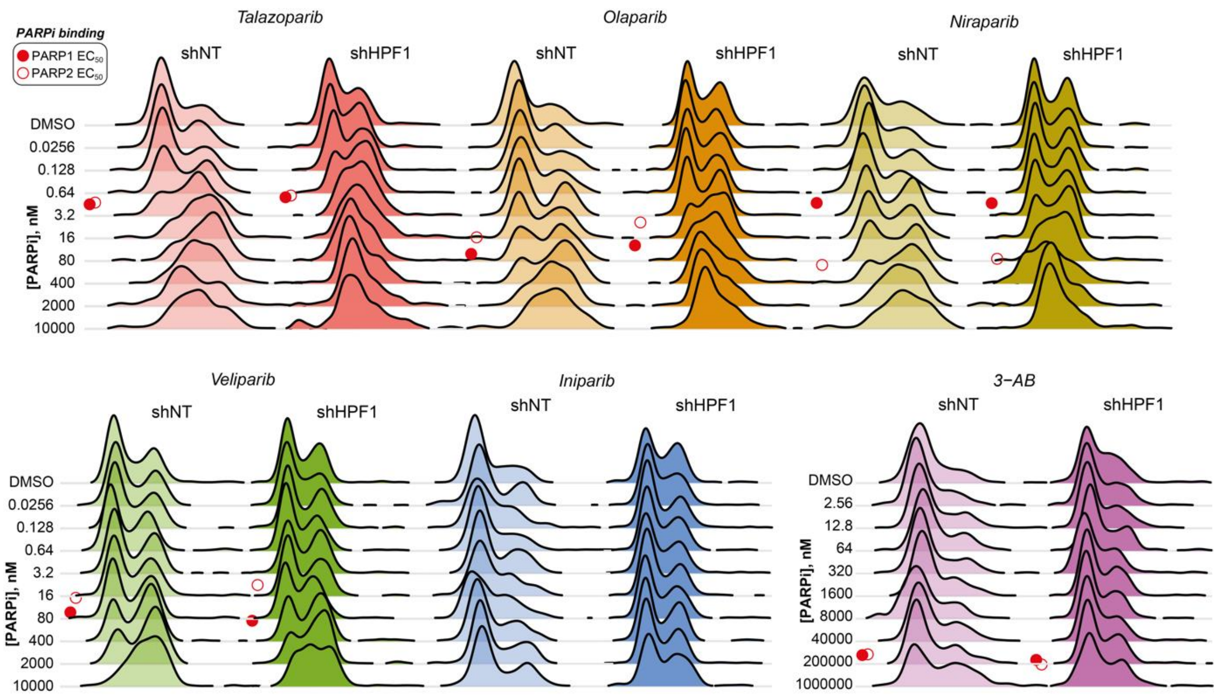


**Supplementary Figure 8. Influence of HPF1 depletion on PARPi cell cycle dynamics.** Representative cell cycle profile dynamics from Hoechst intensity of shNT or shHPF1 cells after 24-hour PARPi dose-response, as shown by Ridgeline plot. 3-AB had a concentration range of 2.56 to 1,000,000 nM, all others – 0.0256 to 10,000 nM. Red circles – approximate stabilization $EC_{50}$ values for PARP1 (closed) or PARP2 (open) biosensors.
